# LARP1 senses free ribosomes to coordinate supply and demand of ribosomal proteins

**DOI:** 10.1101/2023.11.01.565189

**Authors:** James A. Saba, Zixuan Huang, Kate L. Schole, Xianwen Ye, Shrey D. Bhatt, Yi Li, Winston Timp, Jingdong Cheng, Rachel Green

## Abstract

Terminal oligopyrimidine motif-containing mRNAs (TOPs) encode all ribosomal proteins in mammals and are regulated to tune ribosome synthesis to cell state. Previous studies implicate LARP1 in 40S- or 80S-ribosome complexes that repress and stabilize TOPs. However, a mechanistic understanding of how LARP1 and TOPs interact with these complexes to coordinate TOP outcomes is lacking. Here, we show that LARP1 senses the cellular supply of ribosomes by directly binding non-translating ribosomal subunits. Cryo-EM structures reveal a previously uncharacterized domain of LARP1 bound to and occluding the 40S mRNA channel. Free cytosolic ribosomes induce sequestration of TOPs in repressed 80S-LARP1-TOP complexes independent of alterations in mTOR signaling. Together, this work demonstrates a general ribosome-sensing function of LARP1 that allows it to tune ribosome protein synthesis to cellular demand.

**One-Sentence Summary:** LARP1 directly binds free ribosomal subunits to repress TOP mRNAs

## Main Text

Terminal oligopyrimidine motif-containing mRNAs (TOPs) begin with a +1 cytidine nucleotide followed by 4-15 pyrimidines and encode all ribosomal proteins in mammals (*1*). For decades, it has been known that TOP translation is acutely regulated to coordinate ribosomal protein synthesis with demand across diverse cellular contexts (*2*–*11*) in a manner wholly dependent on the TOP motif at the 5’-end (*12*–*14*). These findings ultimately led to the identification of La-related protein 1 (LARP1) – a multi-domain evolutionarily-conserved RNA binding protein (*15*) – as a key regulator that directly binds to the TOP motif (*16*–*19*).

LARP1 binds TOPs under conditions of mammalian target of rapamycin (mTOR) inhibition, leading to translational repression, as well as stabilization and polyA-tail lengthening of the mRNA (*16*, *19*–*22*). Despite some literature suggesting mTOR components could be dispensable for TOP control (*23*–*25*), mTOR is thought of as the quintessential regulator of TOP translation (*16*, *26*–*28*) in part through direct phosphorylation of LARP1 (*19*, *29*–*33*).

Interestingly, the LARP1-TOP complex itself is thought to be associated with ribosomes as LARP1 and TOPs sediment with 40S ribosome complexes (TOP-40S) during normal growth conditions (*34*) or 80S ribosome complexes (TOP-80S) during mTOR inhibition (*21*, *31*, *35*). One recent report argued that the TOP-80S reflects single-ribosome (monosomal) translation that protects TOPs and permits their rapid reactivation (*35*). A recent CLIP-seq study defined contacts between LARP1 and 18S rRNA of the 40S ribosomal subunit near the mRNA channel (*36*). Despite these mechanistic insights, the function, regulation, and biophysical nature of the interaction of LARP1 and TOPs with 40S and 80S ribosomes is not entirely clear.

Here, we define the association of the LARP1-TOP complex with ribosomes. Through biochemical and structural analysis, we demonstrate that the TOP-80S comprises non-translating, weakly-associated 40S and 60S subunits that bind directly to LARP1 and not to the TOP. We show that free ribosomal subunits bind directly to LARP1 and repress TOPs independent of changes in mTOR signaling previously thought to be fundamental to TOP regulation. Our observations provide molecular insights into how LARP1 senses free ribosomes to tune the synthesis of ribosomal proteins to cellular demand.

### A system to study TOP association with 40S and 80S ribosomes

To study the TOP-80S, we treated cells with either DMSO (control) or the mTOR inhibitor, Torin1 (*37*), fractionated cell lysates across a sucrose gradient and performed nanopore sequencing of mRNAs isolated from each fraction (Fig. 1A). Normalization to spike-in mRNAs enabled us to quantify the percent distribution of each mRNA species in each fraction across the gradient. In DMSO, TOPs sediment in a bimodal distribution (Fig. 1B): one population sediments in sub-polysomal fractions, representing translationally-repressed mRNAs associated with 40S subunits (*4*, *21*, *34*), and the other population sediments in polysomal fractions, representing highly-translated mRNAs. In contrast, non-TOPs are more uniformly distributed across polysomal fractions, indicative of active translation. Upon Torin1 treatment, TOPs redistribute *en masse* to the 80S fraction (Fig. 1B). In contrast, non-TOPs shift to lighter fractions but remain distributed across the gradient, consistent with a modest global decrease in their translation upon Torin1 treatment (Fig. 1B). These data agree with previous reports showing stronger translational repression of TOPs compared to non-TOPs with Torin1 treatment (*20*, *28*). Importantly, qPCR for individual TOPs recapitulated the same redistribution from the 40S and polysomal fractions to the 80S fraction upon Torin1 treatment (Fig. 1C and fig. S1A). These data therefore define a robust system for interrogating the sedimentation of TOPs with 40S and 80S ribosomes.

**Fig. 1.**
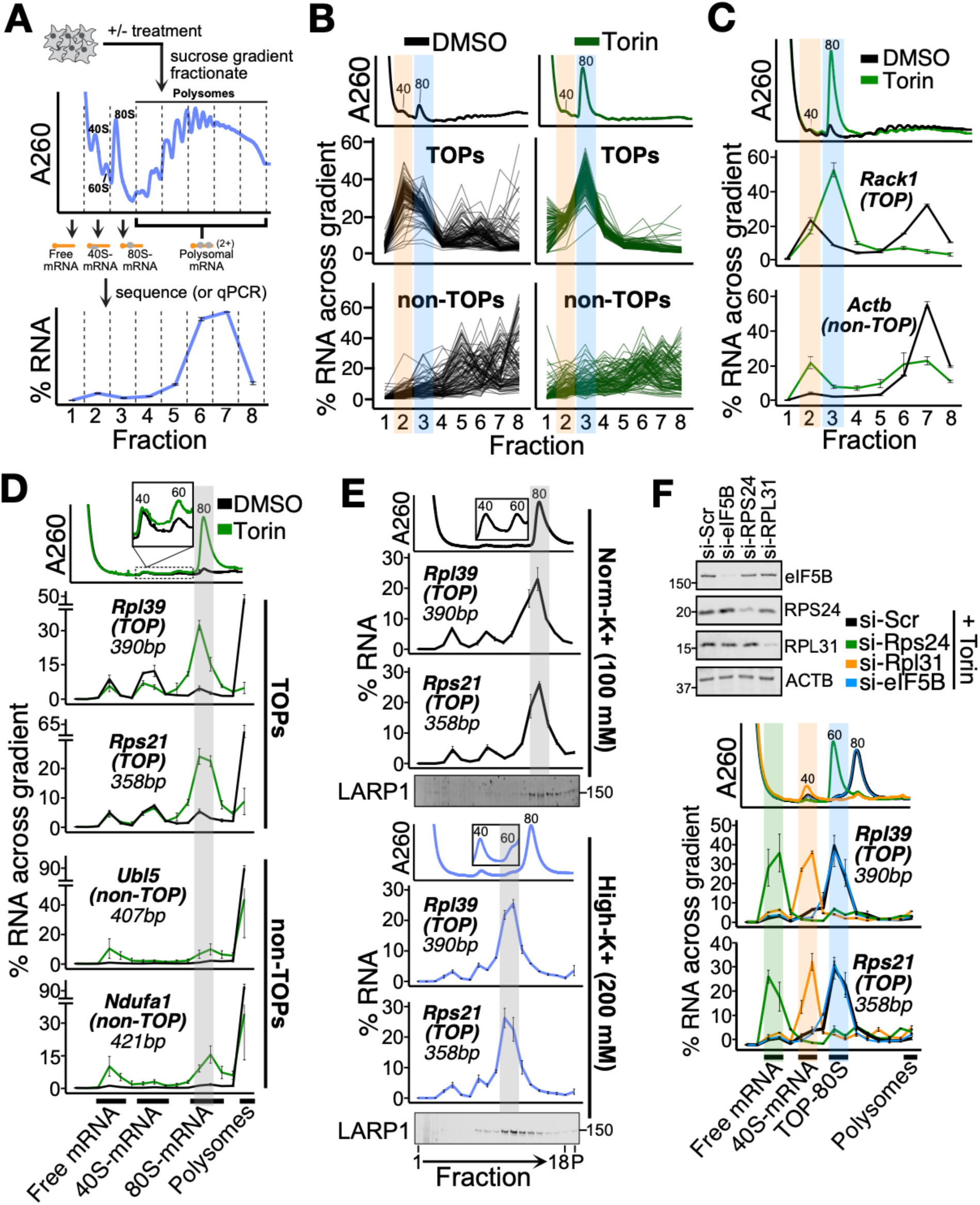
Characterization of the TOP-80S complex. (**A**) Schematic of experimental setup. Cell lysates were separated by ultracentrifugation through a sucrose gradient and fractionated. The amount of each RNA species in each fraction is presented as a percentage of the whole across the gradient as shown here for a representative mRNA. (**B**) U266B1 cells were treated with DMSO or 300 nM Torin1 for 1 h, lysed, and processed as shown in (A) followed by PCR-cDNA nanopore sequencing. A260 traces for each treatment are shown in upper panels. Each line represents the fractional distribution of a single mRNA across the gradient. Top: 96 annotated TOPs are plotted +/- Torin1. Bottom: A random sample of 96 non-TOPs are plotted +/- Torin1. Orange and blue highlights correspond to the TOP-40S and TOP-80S, respectively. (**C**) HEK293T cells were treated with DMSO or 300 nM Torin1 for 1 h and processed as shown in (A) followed by qPCR against genes of interest. Orange and blue highlights correspond to the TOP-40S and TOP-80S, respectively. (**D**) HEK293T cells were treated with DMSO or 300 nM Torin1 for 1 h and lysates fractionated on 15-35% sucrose gradients containing 100 mM KOAc followed by qPCR against genes of interest. Gray highlight corresponds to the TOP-80S. (**E**) HEK293T cells were treated with 300 nM Torin1 for 1 h and lysates fractionated along 15-35% sucrose gradients containing either 100 mM KOAc (Norm-K^+^) or 200 mM KOAc (High-K^+^) followed by qPCR against genes of interest. Gray highlight corresponds to the TOP-80S. Western blots against LARP1 from the same samples are presented below qPCR traces. (**F**) HEK293T cells were treated with siRNAs targeting scrambled (si-Scr), Rps24, Rpl31, or eIF5B followed by 300 nM Torin1 for 1 h. Western blots demonstrating knockdown efficiency are shown. Lysates were fractionated along 15-35% sucrose gradients containing 200 mM KOAc followed by qPCR against genes of interest. Green, orange, and blue highlights correspond to free mRNA, TOP-40S, and TOP-80S, respectively. For (C-F), data are shown from one experiment representative of two biological replicates. For qPCR plots, error bars reflect the SD of 2-4 technical replicates from one experiment.

### TOP-80S ribosomes are weakly-associated and non-translating

To better resolve the messenger ribonucleoprotein (mRNP) complexes in the sub-polysomal fractions, we ultracentrifuged cellular lysates over lower percentage sucrose gradients, fractionated finely, and probed for size-matched, endogenous mRNAs whose annotated size of < 500 bp should minimally affect the sedimentation of mRNPs. Hereafter we refer to these as “spread” gradients. This protocol allowed us to identify three distinct populations of TOPs which sediment in the sub-polysomal region of the gradient: 1) Free mRNA, 2) TOP-40S and 3) TOP-80S (Fig. 1D). We also extracted the material that sedimented to the bottom of the ultracentrifuge tube. This allows us to determine the proportion of each mRNA species in polysomes and therefore account for the full cytoplasmic expression of a given mRNA species.

As seen previously, after Torin1 treatment TOPs (Rpl39 and Rps21) exit polysomes and sediment with 80S monosomes (Fig. 1D). In contrast, non-TOPs (Ubl5 and Ndufa1) are more resistant to Torin1 treatment: a substantial portion remains in polysomes and a moderate portion sediments with the free mRNA or 80S peaks. In accordance with previous studies (*21*, *35*), the TOP-80S fails to form in LARP1-KO cells upon Torin1 treatment, confirming that LARP1 is a critical component of this complex (fig. S1B).

Because of the high resolution afforded by these spread gradients, we were able to resolve a subtle leftward shift of TOPs in the 80S region compared to non-TOPs (Fig. 1D). We wondered whether the ribosome species associating with TOPs might be qualitatively different from an elongating 80S monosome. We treated cells with Torin1 and then compared the sedimentation of TOPs in gradients containing 100 mM (“norm-K^+^”) or 200 mM (“high-K^+^”) KOAc. While TOPs sediment with the leftmost edge of the monosome peak in norm-K^+^ gradients, they sediment even further left (closer to the 60S distribution) in high-K^+^ gradients (Fig. 1E). This finding is reminiscent of early work showing that high KOAc causes vacant 40S and 60S couples (but not mRNA-engaged ribosomes) to split as they travel through the gradient, leading to intermediate migrations of the dissociating 40S and 60S components (*38*–*40*) (see supplementary text on “TOP-80S shift in high-K^+^ gradients”). Importantly, LARP1 protein undergoes the same redistribution in high-K^+^ gradients, landing in the TOP-80S left-shifted peak (Fig. 1E). Titrating KOAc even higher to 500 mM causes TOPs to sediment still further left, likely reflecting earlier dissociation in the gradient (fig. S1C). Based on these data, we propose that the TOP-80S contains vacant 40S and 60S ribosomal subunits which are not directly engaged in translating the bound mRNA.

To directly evaluate whether both 40S and 60S subunits are present in the TOP-80S, we treated cells with Torin1 following siRNA-mediated knockdown of RPS24 (a 40S protein) or RPL31 (a 60S protein). These knockdowns impacted the overall abundance of 40S and 60S subunits as previously observed for other ribosomal protein knockdowns (*34*). Importantly, si-RPS24 resulted in migration of TOPs entirely in the free mRNA fraction (with no ribosomal subunits bound) while si-RPL31 resulted in migration of TOPs entirely in the TOP-40S fraction (Fig. 1F). These data confirm that the TOP-80S contains both a 40S and a 60S subunit. Importantly, these data also demonstrate that the 40S joins first and directly to the TOP because knockdown of the 40S prevents both subunits from joining while knockdown of the 60S prevents only 60S joining.

To test our model that the TOP-80S does not contain actively elongating monosomes, we knocked down eIF5B, an initiation factor which mediates 60S joining at AUG start codons (*41*). In untreated cells, eIF5B knockdown caused an increase in the monosome peak confirming that the knockdown was sufficient to globally inhibit translation initiation (fig. S1D). Strikingly, eIF5B knockdown did not impact the migration or abundance of the TOP-80S under Torin1 treatment (Fig. 1F). These data demonstrate that 60S joining to the TOP-80S is not mediated by canonical mechanisms (eIF5B) at an AUG start codon. Instead, consistent with the high-K^+^ data (Fig. 1E), these data suggest that the TOP-80S is composed of 40S and 60S subunits bound to one another by more general inter-subunit interactions independent of translation initiation. Surprisingly, we also noticed that eIF5B knockdown strongly induced formation of the TOP-80S in untreated cells, phenocopying what we had observed with Torin1 (fig. S1D). This provides further evidence that 60S joining to LARP1-TOP complexes is not occurring by canonical mechanisms. Taken together, these data demonstrate that the TOP-80S contains a TOP mRNA complexed with LARP1, a 40S subunit and a weakly-associated 60S subunit, and is likely non-translating.

### Cryo-EM structures reveal LARP1 bound in the mRNA channel of the 40S subunit

Intrigued by the association of LARP1 and TOPs with 40S subunits, we solved a single-particle cryo-EM structure of mature human 40S ribosome complexes with LARP1 at 3.2 Å resolution, obtained from an *in vivo* immunoprecipitation using PYM1 (a known ribosome-associated factor (*42*)) as purification bait (Figs. 2A-2B, figs. S2A, S3A, S4A, and table S1). The resolved density maps allowed us to construct a *de novo* model of the 40S ribosome and unambiguously assign and fit AlphaFold (*43*) predicted models for amino acids 660-724 of the long-isoform annotation of LARP1 (ENSEMBL LARP1-204) (*44*) (see supplementary texts on “Validation of LARP1 structure from PYM1 IP” and “LARP1 isoforms”). This assignment of LARP1 was further confirmed by a second cryo-EM structure which we obtained using LARP1 as purification bait (Fig. 2C and figs. S2B, S4B). Based on these two structures we were able to define a previously uncharacterized domain of LARP1 spanning residues 650-730 between the La (polyA-binding (*45*–*47*)) and DM15 (m^7^G-TOP-binding (*17*, *18*)) domains, which we call the <u>R</u>ibosome <u>B</u>inding <u>R</u>egion (RBR; Fig. 2A). Importantly, while LARP1 is found in an identical position in both structures, the superior structure obtained from the PYM1 sample is used for discussion in the remainder of the paper.

**Fig. 2.**
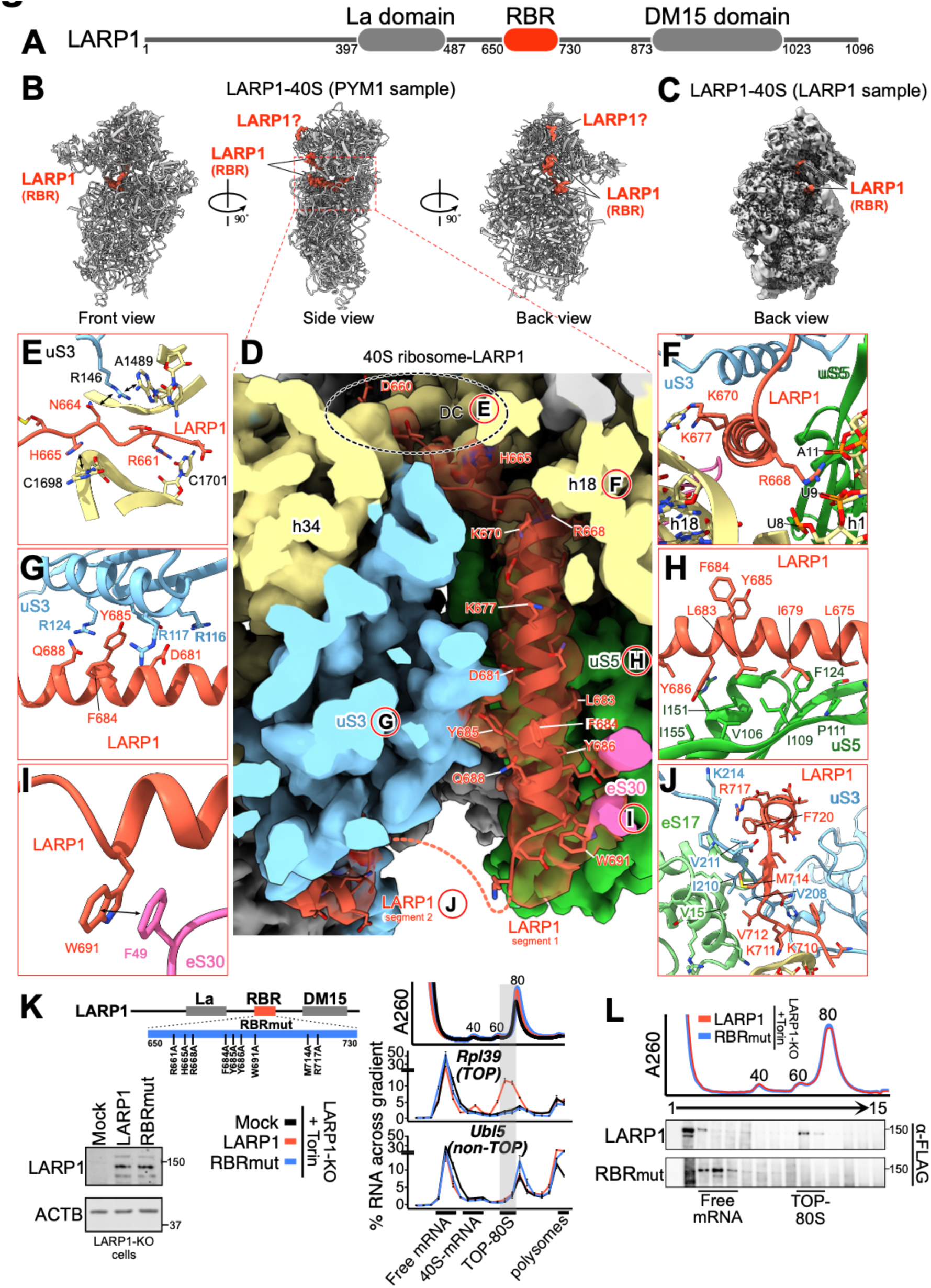
Cryo-EM analysis of the interaction between LARP1 and the 40S ribosome. (**A**) Schematic showing the domain arrangement of LARP1 protein. (**B**) Three different views of cryo-EM structure of the human LARP1-40S ribosome complex (from PYM1 sample). (**C**) Back view of cryo-EM structure of the LARP1-40S ribosome complex (from LARP1 sample). (**D**) Zoom-in of the RBR of LARP1 bound within the mRNA channel of the 40S ribosomal subunit. References to specific figure panels are annotated with the corresponding letter circled in red. (**E**) LARP1 residues 660-665 bind to the decoding center of the 40S ribosome. Key residues interacting with uS3 (blue) and 18S rRNA (yellow) are shown. (**F-I**) Detailed schematics of LARP1 residues 666-694; key residues interacting with 18S rRNA helices 1 and 18 (yellow, F), uS3 (blue, G), uS5 (green, H) and eS30 (pink, I) are shown. (**J**) LARP1 residues 708-724 interacting with 18S rRNA (yellow), uS3 (blue) and eS17 (light green) are shown. For (D-J), residues and nucleobases are annotated and stacking interactions are indicated by black bidirectional arrows. (**K**) LARP1-KO cells (HEK293T) were transfected with plasmids expressing either FLAG peptide alone (Mock), LARP1, or LARP1-RBRmut followed by treatment with 300 nM Torin1 for 1 h. The domain map detailing mutated residues in the RBRmut (top left) and western blots showing LARP1 expression (bottom left) are shown. Lysates were fractionated along 15-35% sucrose gradients containing 200 mM KOAc followed by qPCR against genes of interest. Gray highlight corresponds to the TOP-80S. Data are shown from a single experiment representative of two biological replicates. For qPCR data, error bars reflect the SD of 2-4 technical replicates from one experiment. (**L**) LARP1-KO cells (HEK293T) were treated as described in (K). Protein was extracted from the gradient fractions and western blotted for FLAG-tagged LARP1.

Only the RBR of LARP1 could be observed in our structure and extends from the inter-subunit side to the solvent side of the 40S subunit through the mRNA entry channel (Fig. 2B-2C and figs. S4A-S4B). The RBR can be subdivided into three segments of defined density separated by short stretches of unresolved density (Fig. 2D and figs. S4C-S4E). The first segment of residues 660-694 traverses from the decoding center (DC) through the mRNA channel (Fig. 2D and fig. S4C). The N-terminus of this segment is located within the DC where residue H665 stacks with base C1698 (C1397 in *E. coli*) of the 18S rRNA which plays an important role during tRNA decoding (Fig. 2E) (*48*). This localization agrees with PAR-CLIP data identifying contacts between LARP1 and nucleotides 1698-1702 of the 18S rRNA (*36*). The subsequent protein helix (residues 667-694) passes through the mRNA channel using basic residues to interact with 18S rRNA helices 1 and 18 (e.g. R668 with h1 and K670 with h18; Fig. 2F). Deeper in the channel the same helix makes hydrophobic contacts with ribosomal proteins uS3, uS5, and eS30: Y685 of LARP1 stacks with R117 and R124 of uS3 (Fig. 2G); Y686, L683 and I679 of LARP1 contact the hydrophobic surface of uS5 (Fig. 2H); and W691 of LARP1 stacks with F49 of eS30 (Fig. 2I). Following this helix, LARP1 becomes disordered and the second segment of the RBR emerges and includes residues 708-724 which make hydrophobic contacts with the C-terminal tail of uS3 and with the N-terminal domain of eS17 (Fig. 2J and fig. S4D). Finally, an unknown density on the surface of RACK1 is plausibly a third segment of the RBR (fig. S4E). However, we were unable to confidently assign this region due to the lack of structural features and low local resolution.

The cryo-EM structure of LARP1 bound to 40S subunits within the DC and mRNA channel immediately suggests that the complex is translationally inactive. Indeed, superimposing the mRNA density from a solved structure of a scanning 48S preinitiation complex (PIC) (*49*) reveals that the LARP1 RBR directly clashes with mRNA density in the channel and would occlude mRNA binding (fig. S5A). We further speculate that LARP1 binding to the DC could hinder the association of eIF1A, a critical initiation factor (fig. S5B) (*50*). Finally, the LARP1 RBR occupies a similar position in the mRNA channel to known translation repressors SARS-CoV-2 NSP1 (*51*) (fig. S5C) and human SERBP1 (*52*) (fig. S5D). These observations suggest these factors may employ a conserved mechanism for translation inhibition.

To validate the LARP1-40S structure we introduced nine alanine point mutations into full-length LARP1 at important contact sites with the 40S subunit (“LARP1-RBRmut”) (Fig. 2K). We transfected plasmids expressing either FLAG peptide alone (Mock), FLAG-tagged wild-type LARP1 (LARP1), or FLAG-tagged LARP1-RBRmut (RBRmut) into LARP1-KO cells and treated with Torin1 to evaluate formation of the TOP-80S. Importantly, both the LARP1 and RBRmut constructs are equally expressed and stable as determined by western blot (Fig. 2K).

While the TOP-40S and TOP-80S complexes readily form in cells expressing exogenous LARP1, these complexes fail to form in cells expressing either Mock or the RBRmut; these data establish that the RBR of LARP1 is required for the formation of TOP-40S and TOP-80S (Fig. 2K). By comparison, the sedimentation profile of a non-TOP mRNA, Ubl5, is largely unperturbed in these different backgrounds (Fig. 2K). Furthermore, while wild-type LARP1 protein sediments in the same distribution as the TOP-80S in Torin1-treated cells, the RBRmut protein sediments with free mRNA. These data suggest that the RBRmut LARP1 protein can still bind mRNAs but can no longer bind ribosomes (Fig. 2L). Collectively, these data show that the TOP-40S and TOP-80S are composed of non-translating ribosomal subunits, wherein the 40S subunit is directly bound to LARP1 and not to the TOP itself.

### LRRC47 is a potential new player in regulating TOP translation

In addition to the LARP1-40S ribosome structure, we obtained a LARP1-LRRC47-40S structure from the same *in vivo* PYM1 immunoprecipitation at 3.4 Å resolution (figs. S2A, S3B, and S4F). We were able to construct a *de novo* model of the 40S ribosome and unambiguously assign and fit the AlphaFold (*43*) predicted models for LRRC47. LRRC47 has an N-terminal leucine rich repeat (LRR) domain and C-terminal B3/B4 tRNA-binding domain (Fig. 3A) and was previously captured on late cytoplasmic pre-40S biogenesis intermediates (*53*). In our LARP1-LRRC47-40S structure, LARP1 is in an identical position as seen for the LARP1-40S structure (figs. S4A, S4B, and S4F) and LRRC47 is positioned identically to what was observed in the pre-40S intermediate state F1 (Fig. 3B) (*53*). The LRR domain of LRRC47 binds h11, h22, and h24 and its B3/B4 tRNA binding domain binds h5, h14, and h44 of the 18S rRNA (Fig. 3B) (*53*). The LRR domain would spatially clash with eIF3C and the B3/B4 tRNA binding domain would spatially clash with ABCE1 (*53*, *54*) (Fig. 3B), immediately suggesting that LRRC47 would interfere with assembly of the 43S pre-initiation complex (PIC). These observations are consistent with the model proposed above wherein LARP1-bound 40S complexes are translationally inactive.

**Fig 3.**
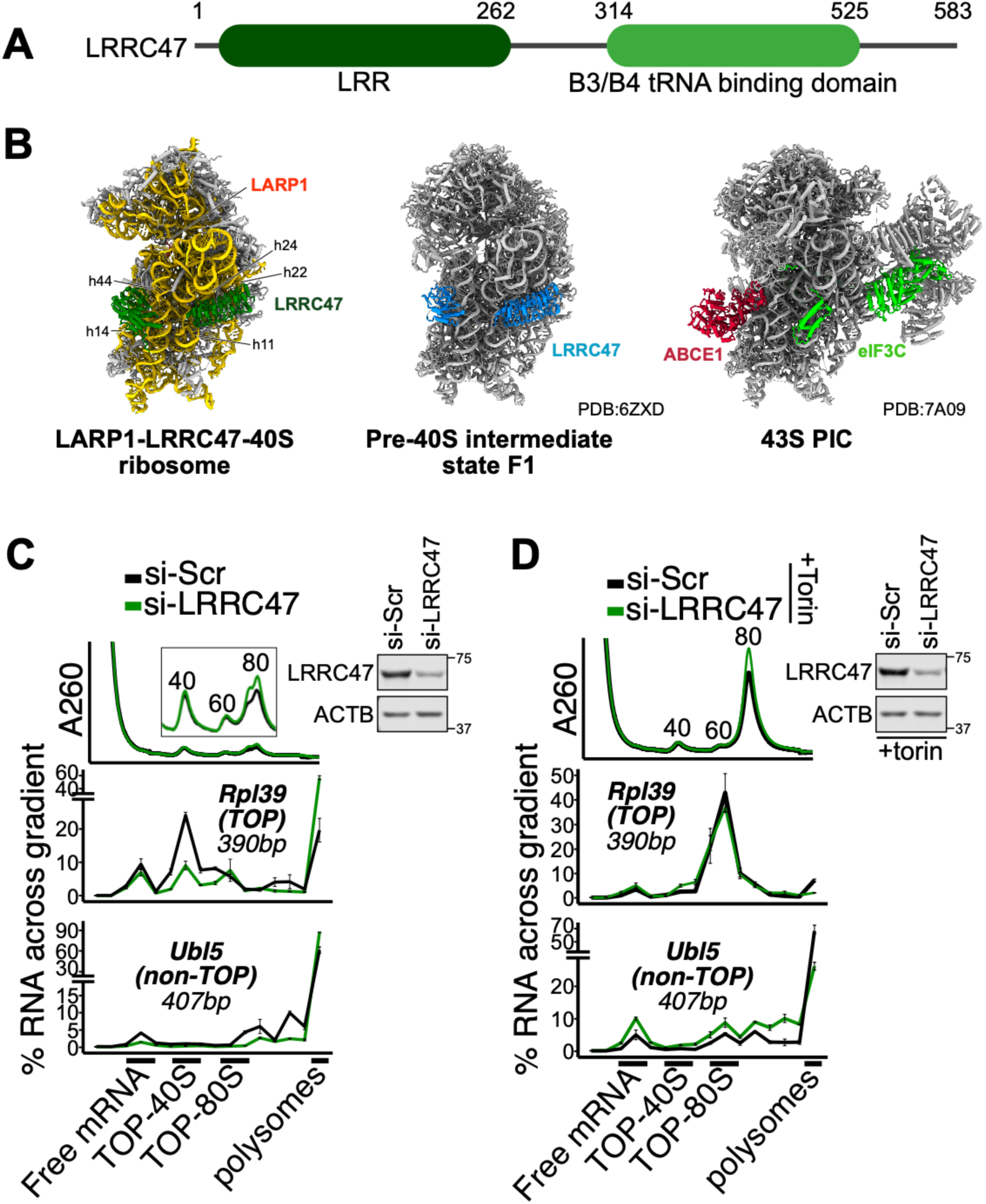
LRRC47 promotes the TOP-40S. (**A**) Domain map of LRRC47 showing LRR (dark green) and B3/B4 tRNA-binding (light green) domains. (**B**) Structural comparison of the LARP1-LRRC47-40S ribosome (left), pre-40S ribosome state F1 (middle) and 43S pre-initiation complex (right). LARP1 (tomato red), LRRC47 (green and blue), ABCE1 (red) and eIF3C (light green) are shown. PIC: pre-initiation complex. (**C**) HEK293T cells were treated with siRNAs targeting scrambled (si-Scr) or LRRC47 and lysates fractionated along 15-35% sucrose gradients containing 200 mM KOAc followed by qPCR against genes of interest. Western blots showing knockdown efficiency are shown. (**D**) Same setup as in (C) except that cells were treated with 300 nM Torin1 for 1 h prior to lysis. For (C-D), data are shown from a single experiment representative of two biological replicates. For qPCR data, error bars reflect the SD of 2-4 technical replicates from one experiment.

While there is limited information on the biological function of LRRC47, it has been hypothesized to act as an anti-association factor as its position on the 40S would preclude 60S joining (*53*). We therefore reasoned that LRRC47 might be involved in forming or stabilizing the TOP-40S (*34*). To test this hypothesis, we evaluated the formation of TOP-40S in cells transfected with scrambled siRNAs or those targeting LRRC47. SiRNA against LRRC47 decreased the abundance of the TOP-40S and increased the proportion of TOPs in polysomes, suggesting that LRRC47 either stabilizes or promotes formation of the TOP-40S under normal growth conditions (Fig. 3C). In contrast, the sedimentation of non-TOPs was largely unperturbed (Fig. 3C). We also evaluated the formation of the TOP-80S in cells treated with Torin1. Knockdown of LRRC47 had no effect on formation of the TOP-80S, suggesting that LRRC47 does not effectively prevent 60S subunit joining to the TOP-40S upon Torin1 treatment (Fig. 3D). Considering that LRRC47 has an unclear role in late-stage 40S biogenesis (*53*), an enticing possibility is that 60S joining under Torin1 conditions kicks LRRC47 off to serve a role in repressing late stages of ribosome biogenesis.

### LARP1 senses free ribosomes to coordinate TOP repression independent of mTOR signaling

The observation that LARP1 binds directly to ribosomes raised the possibility that LARP1 might sense free ribosomes within the cell and couple this function to tuning the translation of TOPs. To test this model, we treated cells with a wide range of conditions broadly known to increase the availability of free ribosomes and probed for formation of the TOP-80S.

Because translational control of TOPs has been extensively studied in the context of mTOR signaling (*16*, *19*–*21*, *26*–*28*, *31*, *35*, *55*–*57*), we began by perturbing downstream components of the mTOR signaling pathway. A major outcome of mTOR inhibition is disruption of the eIF4E-eIF4G interaction that is critical to translation initiation (*58*). As expected, siRNA-mediated knockdown of either eIF4E or eIF4G led to a global decrease in translation initiation as evidenced by an increase in the monosome peak and decrease in polysomes (Fig. 4A).

**Fig. 4.**
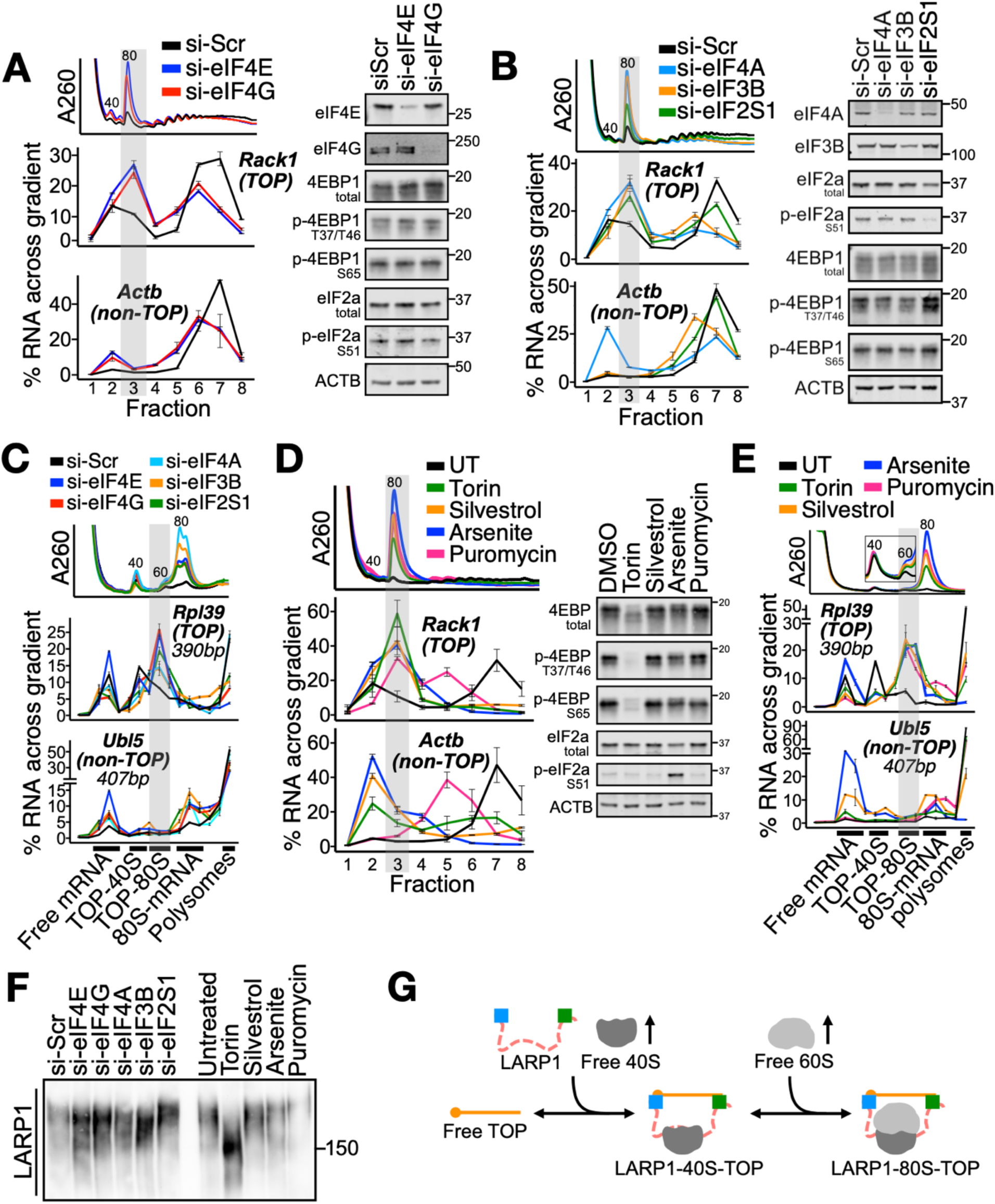
TOPs are repressed in 80S complexes independent of changes in mTOR signaling. **(A)** HEK293T cells were treated with siRNAs targeting scrambled (si-Scr), eIF4E, or eIF4G and lysates fractionated along 10-50% sucrose gradients followed by qPCR against genes of interest. Western blots showing knockdown efficiency and for proteins of interest are shown. (**B**) Identical to (A) except with siRNAs targeting scrambled (si-Scr), eIF4A1/2, eIF3B, or eIF2S1. (**C**) HEK293T cells were treated with siRNAs targeting scrambled (si-Scr), eIF4E, eIF4G, eIF4A1/2, eIF3B, or eIF2S1 and lysates fractionated along 15-35% sucrose gradients containing 200 mM KOAc followed by qPCR against genes of interest. (**D**) Identical to (A) except cells were treated with Torin1 (300 nM, 1 h), Silvestrol (30 nM, 1 h), Sodium arsenite (Arsenite; 100 µM, 1 h), or Puromycin (250 µM, 30 min). (**E**) Identical to (C) except cells were treated with Torin1 (300 nM, 1 h), Silvestrol (30 nM, 1 h), Sodium arsenite (Arsenite; 100 µM, 1 h), or Puromycin (250 µM, 30 min). For (A-E), gray highlights correspond to the TOP-80S. Data are shown from one experiment representative of two biological replicates (A, B, D) or one biological replicate (C, E). For qPCR data, error bars reflect the SD of 2-4 technical replicates from one experiment. (**F**) Phos-tag gel for LARP1 following identical treatments to those described throughout Figure 4. Data are shown from one experiment representative of two biological replicates. (**G**) Model: LARP1 senses increases in free 40S and 60S ribosomal subunits within the cell which induces binding and repression of TOPs. Ribosomes bind directly to the LARP1 RBR which is located between the DM15 domain (blue) and La domain (green).

Importantly, neither condition led to an observable change in eIF4E-binding protein 1 (4EBP1) phosphorylation (a canonical readout of mTOR signaling (*37*)) or eIF2S1/eIF2*α* phosphorylation (a canonical readout of the integrated stress response (ISR) (*59*)) (Fig. 4A). Nonetheless, both conditions led to a redistribution of TOPs, but not non-TOPs, to the 80S fraction by qPCR, indicating formation of the TOP-80S (Fig. 4A and fig. S6A). Furthermore, the distribution of LARP1 protein across the polysome profile revealed increased sedimentation with the 80S fraction upon eIF4E knockdown, consistent with LARP1 involvement in the TOP-80S (fig. S6B). These data demonstrate that the sequestration of TOPs in the TOP-80S can be decoupled from changes in mTOR signaling.

Since eIF4E and eIF4G knockdown should both reduce association of eIF4E with the m^7^G-cap of mRNAs (*60*, *61*), it seemed plausible that these knockdowns might simply allow LARP1 to better compete with eIF4E for the m^7^G-cap of TOPs. We therefore wondered whether formation of the TOP-80S might simply reflect increased accessibility or a more general phenomenon caused by increased free ribosomes. To this end, we next depleted diverse components of the translation initiation machinery, but in this case those not directly affecting cap-binding by eIF4E. As expected, knockdown of eIF4A1/eIF4A2 (dual knockdown), eIF3B, or eIF2S1/eIF2*α* all substantially increased 80S abundance, confirming that global translation initiation was decreased (Fig. 4B). Again, we observed little-to-no decrease in mTOR signaling in any of these conditions. Interestingly, we did note modest upregulation of mTOR activity with siRNA against eIF2S1/eIF2 that likely results from negative crosstalk between the eIF2 and mTOR signaling pathways (Fig. 4B) (*62*, *63*). Remarkably, as with eIF4E and eIF4G depletion, treatment with each siRNA led to the collective redistribution of TOPs, but not non-TOPs, to the 80S fraction (Fig. 4B and fig. S7A), suggesting that increased availability of free ribosomes is sufficient to induce formation of the TOP-80S.

To formally establish the identity of the TOP-80S in these conditions, we repeated the initiation factor knockdowns in spread, high-K^+^ gradients. Importantly, in all conditions, the TOP-80S sediments distinctly upstream of the canonical 80S (Fig. 4C). These data indicate that the TOP-80S under all of these conditions corresponds to non-translating, loosely-associated 40S and 60S subunits which are bound to LARP1 and not to the TOP itself. In contrast, all knockdowns induced a small but appreciable proportion of non-TOP mRNA to sediment with the canonical 80S, providing a nice marker for non-TOP mRNAs which are still translated by a single, elongating monosome (Fig. 4C).

As a final test of the hypothesis that LARP1 senses free ribosomes, we used pharmacological inhibitors to stress cells and acutely increase the availability of free ribosomes. In addition to Torin1 as a positive control, we treated cells with silvestrol (an eIF4A inhibitor (*64*)), sodium arsenite (an inducer of eIF2S1/eIF2*α*-phosphorylation and the ISR (*65*)), or puromycin (a tyrosyl-tRNA mimetic which releases nascent peptides and triggers dissociation of ribosome subunits from mRNAs (*66*)). As expected, acute treatments with all compounds (for 1 hour or less) resulted in a marked increase in monosomes (Fig. 4D). While Torin1 induced an almost complete loss of 4EBP1 phosphorylation at T37/T46 and S65, there was no observable change in mTOR signaling detected with any other agents (Fig. 4D). As expected, sodium arsenite potently activated the ISR, as observable by increased eIF2S1/eIF2*α* phosphorylation and modest translational activation of ATF4 mRNA by qPCR (Fig. 4D and fig. S7B). As with the initiation factor knockdown experiments, treatment with each of these pharmacological agents led to a redistribution of TOPs, but not non-TOPs, to the 80S fraction (Fig. 4D and fig. S7B). Further experiments again showed that the TOP-80S sedimented distinctly upstream of the translating 80S in spread, high-K^+^ gradients (Fig. 4E). Collectively, these data demonstrate that LARP1-mediated translational repression of the TOPs occurs dynamically in response to acute increases in the availability of free ribosomes.

A prevailing model implicates LARP1 phosphorylation at multiple residues as a key mediator of TOP regulation (*19*, *25*, *29*–*33*). Given that none of the pharmacological agents impacted mTOR signaling to 4EBP1, we wondered whether LARP1 phosphorylation status might also be unaffected. We queried LARP1 phosphorylation using Phos-tag gels in which phosphorylated proteins run more slowly than non-phosphorylated proteins. Only Torin1 induced an obvious downshift in the migration pattern of LARP1 by Phos-tag gel while all other treatments that also lead to the formation of the TOP-80S did not lead to an obvious downshift (Fig. 4F). These data indicate that phosphorylation of LARP1 is not appreciably decreased under these regimes and collectively argue that phosphorylated LARP1 can recruit ribosomes and TOPs to repress their translation.

## Discussion

We have defined molecular features of interactions between LARP1, TOPs and 40S and 80S ribosomes. Our biochemistry and structural approaches identified the RBR of LARP1 which directly binds 40S subunits in a manner that prevents mRNA binding and thus translation. In order to form the TOP-80S, the RBR binds a 40S subunit while the DM15 domain simultaneously binds a TOP, and the 60S joins through normal subunit interface interactions that are salt-sensitive. This view is supported by the observation that mutations in either the RBR or DM15 domains preclude TOP-80S formation (our data and (*35*)). Through these coordinated binding events, we propose that LARP1 can directly sense free ribosomes and tune the repression of TOPs accordingly (Model Fig. 4G).

LARP1 exists at an abundance of ∼10^5^ copies per mammalian cell (*67*), roughly stoichiometric with mRNA (*68*) but at least an order of magnitude less abundant than ribosomes (*69*). As such, this sensing mechanism seems well-suited to coordination of TOP repression rather than to sequestration of inactive ribosomes. Importantly, our data show that LARP1-ribosome sensing can occur independent of observable phosphorylation changes in 4EBP1 and LARP1. We acknowledge, however, that signaling downstream of mTOR and other kinases likely provides additional layers of regulation that determine how effectively these repressive complexes form (*19*, *25*, *29*–*33*). For example, it is possible that in certain cell types or regimes, 4EBP1 phosphorylation may play more critical roles in providing access to the TOP m^7^G-cap (*28*, *57*).

While our data do not allow us to assign an order of events for complex formation, we are inclined toward a model in which free 40S subunits first bind LARP1 and this induces binding to TOPs. An exciting possibility would be if ribosome binding induces a conformational change that increases DM15 affinity for the TOP motif. Importantly, our data also do not allow us to claim an essential role for 60S joining to form the TOP-80S, although formation of this complex is robust and ubiquitous in all conditions tested. Formation of this complex may simply reflect the binding equilibrium in the cell that occurs in the presence of an excess of both 40S and 60S subunits. Alternatively, given the established role of LARP1 in stabilizing TOPs (*16*, *34*, *57*, *70*), 60S joining could drive stabilization by mass action (Fig. 4G), allowing for more TOPs to be recruited into the most stable state.

The ribosome-sensing function of LARP1 uncovered here seems likely to be implicated in cell states in which free ribosome concentrations are altered or dynamically regulated such as cell differentiation (*71*), changes in cell size (*72*, *73*), and disease (*74*, *75*). Importantly, LARP1 expression is elevated and associated with adverse prognosis in several cancers including cervical (*76*), non-small cell lung (*76*, *77*), hepatocellular (*78*), colorectal (*79*), and ovarian cancers (*80*). Considering the varying demands on ribosome number in each of these diseases and cell states, LARP1-ribosome sensing could serve as a general mechanism to tune synthesis of ribosomal proteins to the availability of free ribosomes across a diverse array of healthy and diseased mammalian cell contexts.

## Supporting information

Data S1. Mass spectrometry results of the PYM1 immunoprecipitation.

## Acknowledgements

We thank members of the Green, Cheng, and Timp Labs for helpful comments and suggestions during preparation of this manuscript. We thank Paul Hook for help with Nanopore sequencing analysis. We thank Carson Thoreen for providing LARP1-KO cell lines (HEK293T) and LARP1 expression plasmids.

## Author contributions

Conceptualization: JAS, JC, RG

Methodology: JAS, ZH, KLS, JC, RG

Investigation: JAS, ZH, KLS, XY, SB, YL

Validation: JAS, ZH, KLS, XY, SB

Formal analysis: JAS, ZH, JC

Visualization: JAS, ZH, XY, JC, RG

Funding acquisition: WT, JC, RG

Writing - original draft: JAS, JC, RG

Writing - review & editing: JAS, ZH, KLS, XY, SB, YL, WT, JC, RG

## Data and materials availability

All study data are included in the article and/or supporting information. Nanopore sequencing data is deposited in the Gene Expression Omnibus (GSE246077). Code for analyzing sequencing data and qPCR data, and for generating plots has been posted on GitHub at repository 2023_Saba_Larp1 (https://github.com/jakesaba/2023_Saba_LARP1). Any additional data, code, or materials used in this article will be made available upon request. Cryo-EM structural data will be deposited in the Electron Microscopy Data Bank (EMDB) and Protein Data Bank (PDB) to be released upon article publication.

## Materials and Methods

### Cell culture

For HEK293T (ATCC CRL-3216) and HEK293/Flp-In/T-Rex (Invitrogen R71007) cell lines, cells were cultured in DMEM (ThermoFisher 11995073) supplemented with 10% FBS (ThermoFisher A3160502). The HEK293/Flp-In/T-Rex were additionally cultured with 1x penicillin/streptomycin (ThermoFisher 10378016). To begin experiments, cells were trypsinized in 0.25% Trypsin-EDTA (ThermoFisher 25200114), pelleted at 350 x g for 4 mins, resuspended in DMEM/FBS, and seeded to the appropriate concentration in tissue culture dishes (Corning). Experiments were started the following day after allowing one overnight for cell attachment. For U266B1 cell lines (ATCC), cells were cultured in RPMI 1640 GlutaMAX media (ThermoFisher 61870127) supplemented with 20% FBS, 1 mM sodium pyruvate (ThermoFisher 11360070), and 5 ng/uL human IL-6 recombinant protein (ThermoFisher PHC0066).

### SiRNA knockdowns

All siRNAs were purchased from Horizon Discovery as ON-TARGETplus SMARTPools according to the following catalog numbers:

Non-targeting (Scramble): D-001810-01-20

RPS24: L-011155-00-0005

RPL31: L-013587-00-0005

EIF5B: L-013331-01-0005

LRRC47: L-010724-01-0005

EIF4E: L-003884-00-0005

EIF4G1: L-019474-00-0005

EIF4A1: L-020178-00-0005

EIF4A2: L-013758-01-0005

EIF3B: L-019196-00-0005

EIF2S1: L-015389-01-0005

RAPTOR: L-004107-00-0005

RICTOR: L-016984-00-0005

LARP1: L-027187-00-0005

EIF4EBP1: L-003005-00-0005

EIF4EBP2: L-018671-00-0005

SMARTPool siRNAs were resuspended to 50 µM in siRNA buffer (10 mM Tris, 60 mM KCl, 0.2 mM MgCl2) and stored as aliquots at −20°C.

SiRNA knockdowns were performed for 48-72 h using Lipofectamine RNAiMAX transfection reagent (ThermoFisher 13778150) according to the manufacturer instructions with some modifications. Cells were seeded one day prior to initiating siRNA treatment and allowed 24 h to attach. For a 10 cm dish containing 10 mL media, 2-5 µL of 50 µM siRNA stock was resuspended in 500 µL OptiMEM reduced serum media (ThermoFisher 51985034) and gently inverted to mix. Simultaneously, 30 µL of RNAiMAX transfection reagent was added to 500 µL OptiMEM and gently inverted to mix. After 5 min incubation at room temperature, the RNAiMAX-OptiMEM mixture was added dropwise to the siRNA-OptiMEM mixture and gently inverted to mix. After 10 min incubation at room temperature, the entire mixture was added dropwise to the cell media and gently swirled to mix (2-5 nM final siRNA concentration per well). The next day, the cell media was changed and the same siRNA treatment was performed. In total, two siRNA transfections were performed for each dish. Cells were incubated for an additional 24-48 h following the second siRNA transfection before lysis.

For siRNA transfections in 6-well dishes, siRNA reagent amounts were decreased to maintain the same concentration in 3-4 mL of media.

### Sucrose gradient fractionation

Cells were seeded at 0.6-3 x 10^6^ cells per dish in 10 cm dishes (Corning CLS430167) and allowed to attach for ∼24 h before starting on the appropriate drug or siRNA treatment. On the day of lysis, cells were replenished with fresh media 2 h prior to lysis. Cells were then started on appropriate drug treatments 20 min - 1 h prior to lysis (Torin1 1 h prior; Silvestrol 1 h prior; Sodium Arsenite 1 h prior; Puromycin 20-30 min prior). At the time of lysis, cells were washed in 5 mL PBS (ThermoFisher 11995065) containing 360 µM emetine dihydrochloride (Millipore Sigma 324693) and 200 µL of sucrose gradient lysis buffer was added dropwise to the plate (<u>sucrose gradient lysis buffer recipe</u>: 50 mM HEPES pH 7.4, 100 mM KOAc, 15 mM Mg(OAc)2, 5% Glycerol, 0.25% (v/v) NP-40 Alternative (Millipore Sigma 492018), 360 μM emetine dihydrochloride, 1 mM TCEP (Gold-Bio TCEP25), 20 U/mL SUPERaseIn RNAse Inhibitor (ThermoFisher AM2696), 1x Halt protease and phosphatase inhibitor cocktail (ThermoFisher 78444), and 8 U/mL TURBO DNase (ThermoFisher AM2239)). Cells were scraped directly from plate in lysis buffer, gently pipetted to homogenize, transferred to ice for 10 min, and clarified by centrifuging at 8000 x g at 4°C for 5 min.

Sucrose buffers were prepared on ice containing 25 mM HEPES pH 7.4, 100 mM KOAc, 5 mM Mg(OAc)2, 180 µM emetine dihydrochloride, 1 mM TCEP, 4 U/mL SUPERaseIn RNAse Inhibitor, and sucrose to the appropriate concentration (<u>non-spread gradients</u>: 10% and 50% sucrose buffers (w/v); <u>spread gradients</u>: 15% and 35% (w/v) sucrose buffers). For high-K^+^ gradients, the KOAc concentration in the sucrose buffers was 200 mM. Six mL of the low-percent sucrose buffer was pipetted to an SW41 ultracentrifuge tube (Seton Scientific 7030) after which six mL of the high-percent sucrose buffer was added to the bottom of the tube using a syringe and cannula. Sucrose gradients were mixed on a Biocomp Gradient Master and placed at 4°C until use on the same day.

Cell lysates were normalized to RNA content using the Qubit RNA Broad Range assay kit (ThermoFisher Q10211) and equal amounts of RNA were layered on top of prepared sucrose gradients. Gradients were ultracentrifuged in a Beckman SW41 swinging bucket rotor for either 75 min (non-spread gradients) or for 300 min (spread gradients). Following ultracentrifugation, gradients were simultaneously fractionated and A260 absorbance measured using a Biocomp Piston Gradient Fractionator per the manufacturer’s instructions. After fractionating, the volume in the bottom of the SW41 tube was collected as a measure of subcellular material which had sedimented to the bottom of the gradient.

### RNA extractions

Whole-cell lysates or sucrose gradient fractions were taken directly to 3x volumes of TRIzol reagent (Invitrogen 15596026) supplemented with 30 pg - 1 ng of eGFP spike-in RNA (TriLink Biotechnologies L-7601-100), vortexed, and frozen at −20°C overnight. The next day, RNA was extracted according to the manufacturer’s instructions with slight modifications. In brief, 0.2 mL of chloroform was added per 1 mL TRIzol reagent, samples vortexed and centrifuged at 16,000 x g at 4°C for 15 min. After centrifuging, the top aqueous layer was taken to a new tube containing an equal volume of isopropanol and 1.5 µL GlycoBlue coprecipitant (ThermoFisher AM9516). Samples were left at −20°C for > 1 h and centrifuged at 21,000 x g for 30 min. The pellet was washed once with 75% ethanol, spun at 21,000 x g for 10 min, supernatant aspirated, and air dried at room temperature. Once dry, the pellet was dissolved in 20-50 µL low TE buffer (10 mM Tris-HCl, 0.1 mM EDTA) and either used immediately or frozen at −80°C.

### Nanopore sequencing across gradients

U266B1 cells were seeded at a density of 7.5 x 10^5^ cells/mL in 55 mL of media (RPMI supplemented with 20% FBS, 1 mM sodium pyruvate, and 5 ng/µL IL-6) in a T225 flask (CytoOne CC7682-4822). The next day cells were treated with either 300 nM Torin1 or an equivalent volume of DMSO (vehicle) for 1 h. After 1 h, cells from two flasks per condition were pooled and spun at 1000 x g for 1 min at room temperature, the media aspirated, and the cell pellet directly lysed in 150 µL of sucrose gradient lysis buffer. Lysates were gently pipetted to homogenize, transferred to ice for 10 min, and clarified by centrifuging at 8000 x g at 4°C for 5 min. Cell lysates were normalized to RNA content using the Qubit RNA Broad Range assay kit (ThermoFisher Q10211) and ∼120 µg of RNA was sedimented along 10-50% sucrose gradients for 75 min and subsequently fractionated to 8 fractions. Thirty pg of eGFP mRNA was spiked into each fraction and RNA was extracted as described in “RNA extractions.” Five µg of extracted RNA from each fraction was subjected to polyA-enrichment using the NEBNext Poly(A) mRNA Magnetic Isolation Module (NEB E7490) according to the manufacturer’s instructions. At this stage, an RNA pico bioanalyzer chip (Agilent 5067-1513) was performed and RNA was confirmed to be of high quality. One ng of polyA-enriched RNA from each sample was subjected to the Nanopore PCR-cDNA Barcoding kit (ONT SQK-PCB109) according to the manufacturer’s instructions with 14 cycles of PCR amplification for all libraries. All libraries were run on a Genomic DNA Screen Tape (Agilent 5067-5366) to confirm that the libraries were of high quality. Barcoded DMSO and Torin1 libraries were separately pooled to 100 fmol in 11 µL of elution buffer and each pool was sequenced on R9.4.1 MinION flowcells (ONT; FLO-MIN106) run on a GridION Mk1 sequencing device (ONT; GRD-MK1) for 72 h with an initial bias voltage of −180 mV. In total, 9 barcoded libraries were sequenced on each flow cell (i.e. DMSO: 8 fractions and whole-cell library).

### Analysis of nanopore sequencing data

Sequencing was basecalled in real-time using the on-device Guppy basecaller (v4.2.3) in the MinKNOW software (v20.10.6) using the “high-accuracy” basecalling model. Multiplexed samples were demultiplexed in real-time, also using the on-device Guppy basecaller with the following key settings: “barcoding_kits=[“SQKPCB109”], trim_barcodes=“off”, require_barcodes_both_ends=“off”, detect_mid_strand_barcodes=“off”, min_score=60.”

Each demultiplexed library was separated into its own fastq file using the grep command at the command line. The reference transcriptome FASTA was downloaded from Gencode (human release 38) using the wget command (http://ftp.ebi.ac.uk/pub/databases/gencode/Gencode_human/release_38/gencode.v38.transcripts.fa.gz) and the sequence for the eGFP spike-in (TriLink L-7601) was manually appended to the reference transcriptome (https://www.trilinkbiotech.com/media/folio3/productattachments/product_insert/egfporf_catno_l-7201_l-7601_l-7701_.txt).

Each library was then mapped to the transcriptome (using minimap2 with the -ax splice option), non-primary alignments excluded (using samtools view with the -F 256 option), and each output BAM file sorted (using samtools sort). Alignments were quantified by counting the number of reads aligning to each transcript within each BAM file (using the uniq -c command at the command line) and raw counts were output as a CSV file. CSV files were subsequently imported into R for further analysis and plotting.

In R, CSV files were joined into a large data frame. NA values were removed and transcripts were filtered for those with at least 80 total reads across all 8 fractions combined (i.e. on average 10 reads or more per fraction). Unmapped reads were removed and transcriptome quantifications were normalized to the eGFP quantification within each sample. Percent RNA across the gradient was calculated as the eGFP-normalized value within each fraction relative to the total for all fractions across the gradient. To collapse transcriptome quantifications to the gene level, the ENSEMBL canonical transcript was then pulled from biomaRt (ENSEMBL version 110) and only the canonical transcript for each gene was kept in the data frame. Plots were constructed in R using the ggplot2 package. All relevant code is published on GitHub at repository 2023_Saba_Larp1 (https://github.com/jakesaba/2023_Saba_LARP1).

### RT-qPCR

Reverse transcription (RT) of extracted RNA (either from whole-cell lysates or sucrose gradient fractions) was performed using the ProtoScript II First Strand cDNA Synthesis Kit (NEB E6560L) according to the manufacturer instructions with some modifications. Six µL of extracted RNA was resuspended in 2 µL of random hexamer primer, heated to 95°C for 2 mins and immediately placed on ice. To each reaction, 10 µL of 2x reaction mix and 2 µL of enzyme mix were added and mixed by pipetting on ice. Reactions (20 uL total) were incubated at 25°C for 5 min, 42°C for 60 min, and 80°C for 5 min in a thermal cycler.

Following RT, cDNA was diluted 1:4 and 4.3 µL of cDNA was mixed with 5 µL iTaq Universal SYBR Green Supermix (BioRad 1725124) and 0.7 µL of 5 µM forward and reverse qPCR primers targeting the gene of interest (333 nM final primer concentration) using the following cycle times: 95°C for 2 min, [95°C for 10 s, 60°C 30 s] × 40 cycles, and then 60°C for 3 min on a QuantStudio 6 RT-PCR system.

#### Analysis

For each experiment, a standard curve containing seven 1:2 dilutions of a stock cDNA was run for each primer set and the relative quantity of cDNA within each qPCR reaction was interpolated from the standard curve. Additionally, qPCR reactions targeting the eGFP spike-in RNA were performed for all sucrose gradient fractions within each experiment to control for differences in RNA extraction efficiency or RT. The quantification for each gene of interest was then normalized to that of the eGFP spike-in. Finally, the percent RNA across the gradient was calculated as a percentage of the eGFP-normalized cDNA value for each fraction relative to the total for all fractions across the gradient for that sample/qPCR target. RT-qPCR reactions were performed in duplicate or triplicate. Plots were constructed in R using the ggplot2 package.

All primer sets used in this study were validated by running an agarose gel of the product and observing a single band at the expected size. The primer sets are displayed below:

EGFP_F: CCCGACAACCACTACCTGAG

EGFP_R: GTCCATGCCGAGAGTGATCC

ACK1_F: GCTGATGGCCAGACTCTGTT

ACK1_R: TTCTAGCGTGTGCCAATGGT

PL11_F: TCCATCATGGCGGATCAAGG

PL11_R: TGTGCAGTGGACAGCAATCT

RPL5_F: CAGCGTATGCACACGAACTG

RPL5_R: ACCTATTGAGAAGCCTGCGG

IF3H_F: TGCTCATTGCAGGCCAGATA

IF3H_R: GGCCATGAAGAGCTTGCCTA

PL39_F: CTGCTCGCCATGTCTTCTCA

PL39_R: CGAATCCACTGGGGAATGGG

PS21_F: AATCGCATCATCGGTGCCAA

PS21_R: CCTGTGACCTTGTCAACCTCG

ACTB_F: ACGTTGCTATCCAGGCTGTG

ACTB_R: GAGGGCATACCCCTCGTAGA

EBP1_F: GGAGTGTCGGAACTCACCTG

EBP1_R: ACTGTGACTCTTCACCGCC

ATF4_F: ATGGGTTCTCCAGCGACAAG

ATF4_R: GGAGAAGGCATCCTCCTTGC N

UFB9_F: GCTTGTTTGATGAGAGCCC N

UFB9_R: AGCACCATTCTGGGACCTT

MYC_F: AGTGGAAAACCAGCAGCCT

MYC_R: TTCTCCTCCTCGTCGCAGTA

UBL5_F: CTCGGGTGAGGAGCTGGT

UBL5_R: TCCTAGCTGGAGCTCGAATC N

UFA1_F: GCGCATCTCTGGAGTTGATCG N

UFA1_R: CAATGTTCTCCAAACCCTTTGACA 18S

RRNA_F: CGAAAGCATTTGCCAAGAAT 18S

RRNA_R: GCATCGTTTATGGTCGGAAC 28S

RRNA_F: CCCAGTGCTCTGAATGTCAA 28S

RRNA_R: AGTGGGAATCTCGTTCATCC

### TCA precipitation of proteins from sucrose gradients

Equal volumes of sucrose gradient fractions were resuspended in 15% trichloroacetic acid (TCA; Millipore Sigma T3699), mixed thoroughly, and stored at −20°C overnight. The following day, samples were centrifuged at 21,000 x g at 4°C for 30 min, aspirated, and pellets washed 2x in 500 µL acetone (centrifuged at 21,000 x g at 4°C for 10 min and aspirated after each wash). After the final wash, pellets were dried at 42°C in a vacuum evaporator for 5 min and resuspended in 2x Laemmli buffer. The total volume of resuspended protein was used for western blotting.

### Western blotting

For whole-cell lysates, clarified lysates were normalized to total protein (by BCA assay; ThermoFisher 23225) or total RNA (by Qubit RNA Broad Range assay kit; ThermoFisher Q10211) in 1X Laemmli buffer. Between 3-10 µg of protein or 1-5 µg of RNA was loaded per lane. For TCA-precipitated samples (from sucrose gradient fractions), the total amount of precipitated protein was loaded in each lane. Samples were boiled at 95°C for 5 min and loaded into 4-12% Criterion XT-Bis-Tris polyacrylamide gels (Bio-Rad 3450125). Gel electrophoresis was performed in 1X MES running buffer (Bio-Rad 1610789) at 150V for ∼1 h. Gels were transferred to nitrocellulose membranes (Bio-Rad 1704271) and blocked in Intercept Blocking Buffer (Licor 927-60001) for 1 h at room temperature. Unless otherwise stated, samples were probed by primary antibody in Intercept Antibody Diluent (Licor 927-65001) at a 1:1000 dilution at 4°C overnight. The next day, membranes were washed 3 x 10 min in TBST at room temperature and hybridized with 800CW goat anti-rabbit IgG secondary antibody (Licor 926-32211) at 1:5000 dilution for 1 h. After 1 h, membranes were washed 3 x 10 min in TBST at room temperature and washed 1 x 10 min in TBS. Blots were imaged using the Licor Odyssey CLx.

All sources and catalog numbers for antibodies used in this study are provided below:

LARP1: Cell Signaling Technology 70180

EIF5B: Proteintech 13527-1-AP

RPS24: Abcam ab196652

RPL31: Raybiotech 144-04089

ACTB: Cell Signaling Technology 4967

FLAG: Cell Signaling Technology 14793 and ABclonal AE024

LRRC47: Proteintech 23217-1-AP

EIF4E: Cell Signaling Technology 9742S

EIF4G: Cell Signaling Technology 2498S

4EBP1: Cell Signaling Technology 9644S

P-4EBP1_T37_T46: Cell Signaling Technology 2855S

P-4EBP1_S65: Cell Signaling Technology 9451S

EIF2ɑ: Cell Signaling Technology 9722S

P-EIF2ɑ_S51: Abcam ab32157

EIF4A: Cell Signaling Technology 2425S

EIF3B: Bethyl Laboratories A301-761A

RAPTOR: Cell Signaling Technology 2280S

RICTOR: Cell Signaling Technology 2140S

RPS3: Abcam ab140688

### Construction of LARP1-RBRmut

pLJC1-LARP1 (WT-LARP1) expresses the C-terminally flag-tagged 1019 amino acid isoform of LARP1 (ENSEMBL LARP1-201) from a CMV promoter as previously described (*19*). The plasmid was linearized by PCR and a gene block introducing 9 alanine point mutations was ordered from IDT and cloned into the linearized plasmid by Gibson assembly (NEB E2611L) according to the manufacturer’s instructions. The point mutations correspond to the following sites of the 1096 amino acid LARP1 isoform (LARP1 ENSEMBL 204; see “supplementary text on “LARP1 isoforms”) (*44*): R661A, H665A, R668A, F684A, Y685A, Y686A, W691A, M714A, R717A. After selection with carbenicillin, constructs were confirmed by whole-plasmid sequencing from single colonies. PcDNA3-FLAG (Addgene 20011) expressing FLAG peptide from a CMV promoter was used as a transfection control in experiments testing LARP1-WT and LARP1-RBRmut constructs.

### Generation of tagged LARP1 and PYM1 cell lines

The human LARP1 and PYM1 genes were amplified from a cDNA library reverse transcribed from SK-HEP1 (ATCC HTB-52) cells and subcloned into a modified pcDNA5/FRT/TO plasmid (Invitrogen V652020), resulting in pcDNA5-LARP1-StFLAG and pcDNA5-PYM1-StFLAG plasmids (C-terminal 2xStrep and 3xFLAG tag). The generation of the stable LARP1 and PYM1 cell lines were adapted from previous work (*81*, *82*). Briefly, HEK 293/Flp-In/T-Rex cells were pre-cultured in a 10 cm dish for one day. At 50% confluence, the cells were co-transfected with 0.5 μg of the corresponding pcDNA5 plasmid and 4.5 μg recombinase plasmid POG44 (Invitrogen V600520). After 48 h, cells were passaged and subjected to a 14-day selection with 200 μg/ml hygromycin B (ThermoFisher 10687010). The selected cell line was validated by Western blotting.

### Native complex preparation and LC-MS

Native ribosome-associated complexes were purified from a stable HEK293/Flp-In/T-Rex cell line as described previously (*81*, *82*). After two days of culture, expression of the bait protein (LARP1 or PYM1) was induced with 1 µg/ml tetracycline (ThermoFisher A39246) for 24 hours. Prior to collection, cells were treated with 10 μg/ml cycloheximide (CHX; Millipore Sigma C1988) for 10 minutes. A total of 50 dishes (15 cm) of cells were collected with a cell scraper, washed with cold 1X PBS buffer, and resuspended in lysis buffer (20 mM HEPES pH 7.4, 100 mM KOAc, 5 mM MgCl2, 1 mM DTT, 10 μg/ml CHX, 0.5 mM NaF, 1 mM Na3V3O4, 1X protease inhibitor mix (Roche 4693132001)). Cells were lysed by 10-15 strokes using a 15 mL douncer. The cell lysate was clarified by centrifugation at 10,000 x g at 4°C for 15 minutes, and the supernatant was incubated with 200 μL FLAG beads (Sigma A2220) in a 50 mL centrifuge tube for 2 hours. After incubation, the beads were transferred to a small chromatography column, (Bio-Rad 7326008) washed once with lysis buffer, and then washed three times with wash buffer (20 mM HEPES pH 7.4, 100 mM KOAc, 5 mM MgCl2, 10 μg/ml CHX). Finally, the protein complex was eluted with 500 μL elution buffer (0.4 mg/ml 3X Flag peptide (Millipore Sigma F4799) in wash buffer) for 45 minutes. The eluate was concentrated using a 100 kDa cut-off concentrator, and the final concentration was determined using a Nanodrop spectrophotometer.

Protein quantification and identification were then performed by label-free protein quantification using LC-MS.

### Electron microscopy and image processing

Purified samples (PYM1 and LARP1 samples, 3.5 μL) were applied to precoated (2 nm) R1.2/1.3 carbon-supported copper grids (Quantifoil), blotted for 4-5 s at 4°C, and plunge-frozen in liquid ethane using an FEI Vitrobot Mark IV. Data were collected on a Titan Krios G4 cryo-electron microscope operating at 300 keV using EPU 2. Cryo-EM data were collected with a pixel size of 1.146 Å/pixel (PYM1 sample) or 0.932 Å/pixel (LARP1 sample) and within a defocus range of −1 to −2.5 μm using a Falcon IV direct electron detector under low dose conditions with a total dose of 50 e-/Å^2^ (PYM1 sample) or 58 e-/Å^2^ (LARP1 sample). Original image stacks were dose-weighted, aligned, summed, and drift-corrected using MotionCor2 (*83*). Contrast transfer function (CTF) parameters and resolutions were estimated for each micrograph using GCTF (*84*). Micrographs with an estimated resolution of less than 5 Å and astigmatism of less than 5% were manually inspected for contamination or carbon breakage.

#### PYM1 sample

A total of 13,347 good micrographs were selected. Automatic particle picking was performed in Gautomatch (https://www2.mrc-lmb.cam.ac.uk/download/gautomatch-053/) without the use of a reference. Picked particles (1,906,164 particles) were extracted in Relion 3.1 (*85*) and then subjected to 2D classification in cryoSPARC (*86*). In the end, 670,975 good particles clearly representing the 40S ribosome were selected for 3D classification in Relion 3.1 (*85*). The detailed sorting scheme is shown in fig. S2A. After two rounds of 3D classification, two classes – LARP1-40S ribosome and LARP1-LRRC47-40S ribosome – displaying strong LARP1 density, were selected and refined to high resolution in Relion (*85*). To obtain the final reconstruction, multibody refinement in Relion was used (*85*). The final maps were post-processed and local resolution filtered using masks that were automatically generated in Relion (*85*). Simultaneously, the final maps were sharpened using DeepEMhancer (*87*).

#### LARP1 sample

A total of 7,643 good micrographs were selected. After automatic particle picking performed in Gautomatch (https://www2.mrc-lmb.cam.ac.uk/download/gautomatch-053/), a total of 518,116 particles were extracted in Relion 3.1 and then subjected to 2D classification in cryoSPARC. Classes (107,728 particles) clearly representing the 40S ribosome were picked together. A non-uniform refinement was performed to obtain the final reconstruction. The final map was also local resolution filtered by cryoSPARC and DeepEMhancer.

### Model building and refinement

In general, we used the human 80S ribosome structures (PDB: 6Z6M) for rigid body fitting in the cryo-EM maps (*52*). Some manual adjustments were made in Coot (*88*), particularly on the 40S head. The Alphafold (*43*) predicted models of LARP1 and LRRC47 were rigid body fitted into their corresponding densities in Coot, then manually adjusted according to the high-resolution maps in Coot. Since the third region of LARP1, which binds on top of RACK1, does not have enough resolution for unambiguous assignment, we omit this model.

The final models were real-space refined with secondary structure restraints using the PHENIX suite (*89*). Final model evaluation was performed using MolProbity (*90*). Maps and models were visualized and figures were generated using ChimeraX (*91*).

### Phos-Tag gel immunoblotting

Cells were seeded to 1-3 x 10^5^ cells per well in a 6-well dish (Corning 3516) and allowed to attach for ∼24 h before starting on the appropriate drug or siRNA treatment. On the day of lysis, cells were replenished with fresh media 2 h prior to lysis. Cells were then started on appropriate drug treatments 20 min - 1 h prior to lysis (Torin1 1 h prior; Silvestrol 1 h prior; Sodium Arsenite 1 h prior; Puromycin 20-30 min prior). At the time of lysis, cells were washed in 1 mL PBS and 100 µL of Phos-Tag lysis buffer was added dropwise to the well (<u>Phos-Tag lysis buffer</u> <u>recipe</u>: RIPA (ThermoFisher 89900), 3X Halt protease and phosphatase inhibitor cocktail (ThermoFisher 78444), ≥40 U/mL benzonase (Millipore Sigma E1014), 1 mM TCEP). Cells were scraped directly from plate in lysis buffer, gently pipetted to homogenize, transferred to ice for 10 min, and clarified by centrifuging at 8000 x g at 4°C for 5 min. Total protein concentrations of clarified lysates were determined by BCA assay (ThermoFisher 23225) and lysates were resuspended to an equal concentration (100-500 ng/µL) in 1X Laemmli buffer and boiled at 95°C for 5 min.

For LARP1 immunoblots, 3-5 µg of total protein was loaded per well and samples were resolved by 5% or 6% SDS-PAGE containing 25 μM Phos-tag reagent (Wako, AAL-107) and 50 μM MnCl2. Gel electrophoresis was performed in 1x Tris/glycine/SDS running buffer (100-125V for 2-4 h) and EDTA-free pre-stained protein marker (Apex Bio F4005) was used as a ladder. Gels were washed twice in 1x transfer buffer (25 mM Tris, 192 mM glycine, 10% v/v methanol) supplemented with 1 mM EDTA followed by two washes in 1x transfer buffer without EDTA (10 min per wash). Gels were transferred to a PVDF membrane (Bio-Rad 1704273) in 1x transfer buffer (35 V, overnight, 4°C). The next day, membranes were blocked in 5% non-fat milk (Santa Cruz Biotechnology sc-2325) w/v in TBST for 1 h and blotted using LARP1 primary antibody at a 1:1000 dilution in 5% milk in TBST at 4°C overnight. The next day, membranes were washed 3 x 10 min in TBST at room temperature and hybridized with mouse anti-rabbit IgG-HRP secondary antibody (sc-2357) at 1:5000 dilution for 1 h. After 1 h, membranes were washed 3 x 10 min in TBST at room temperature and washed 1 x 10 min in TBS. Blots were developed using SuperSignal West Femto Maximum Sensitivity Substrate (ThermoFisher 34095) using a ChemiDoc imaging system (Bio-Rad).

## Supplementary Text

### TOP-80S shift in high-K^+^ gradients

Previous studies have shown that vacant 40S and 60S subunit couples are held together more weakly than ribosomes actively engaged with mRNAs, peptidyl-tRNAs or initiation complexes (*38*–*40*). As a result, these “vacant couples” are susceptible to splitting by high hydrostatic pressure as they travel through sucrose gradients. Hydrostatic pressure follows the following formula:

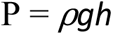

where P = hydrostatic pressure, *⍴* = fluid density, g = acceleration due to gravity, and h = fluid depth. From this formula it becomes clear that as depth in the gradient increases, the hydrostatic pressure proportionally increases. In high-K^+^ gradients, the high potassium concentration further weakens the general intersubunit interactions of vacant couples. As a result, they initially sediment as an 80S ribosome. Once they travel deep enough in the gradient such that the hydrostatic pressure overcomes the force of intersubunit attraction between the 40S and 60S subunits, they split and then begin sedimenting more slowly as independent 40S and 60S subunits. Ultimately this combination of initial migration as an 80S followed by migration as individual subunits causes them and the associated TOP mRNA to sediment in between a normal 40S and 80S distribution.

### Validation of LARP1 structure from PYM1 IP

We were confident in our LARP1 density assignment from the PYM1 immunoprecipitation based on the following: 1) LARP1 was one of the top hits in our semi-quantitative mass spectrometry results of the PYM1 sample (Data S1); 2) The density map has sufficient resolution to resolve all the bulky side-chain information, which nicely fits amino acids 660-724 of LARP1 (figs. S4C-D); and 3) The density map fits the AlphaFold predicted model of LARP1 well with only minor adjustments.

### LARP1 isoforms

While the 1019 amino acid isoform of LARP1 (ENSEMBL LARP1-201) has been referenced extensively in past literature (*16*, *18*, *19*, *31*, *33*, *46*), the 1096 amino acid isoform (ENSEMBL LARP1-204) appears to be the major isoform expressed in most cell types (*44*). Importantly, the two constructs differ only at their N-termini (amino acids 1-144 of the LARP1-204 construct), are identical at characterized domains including the La, RBR, and DM15 domains, and are thought to be functionally equivalent. We used the 1019 amino acid isoform (LARP1-201) for expression of wild-type LARP1 and LARP1-RBRmut constructs (see “Construction of LARP1-RBRmut” in materials and methods). However, all of our annotations and numbering throughout the manuscript correspond to the 1096 amino acid isoform (LARP1-204) to maintain consistency with the isoform that appears to be majorly expressed in human cells (*44*).

---

**Fig. S1.**
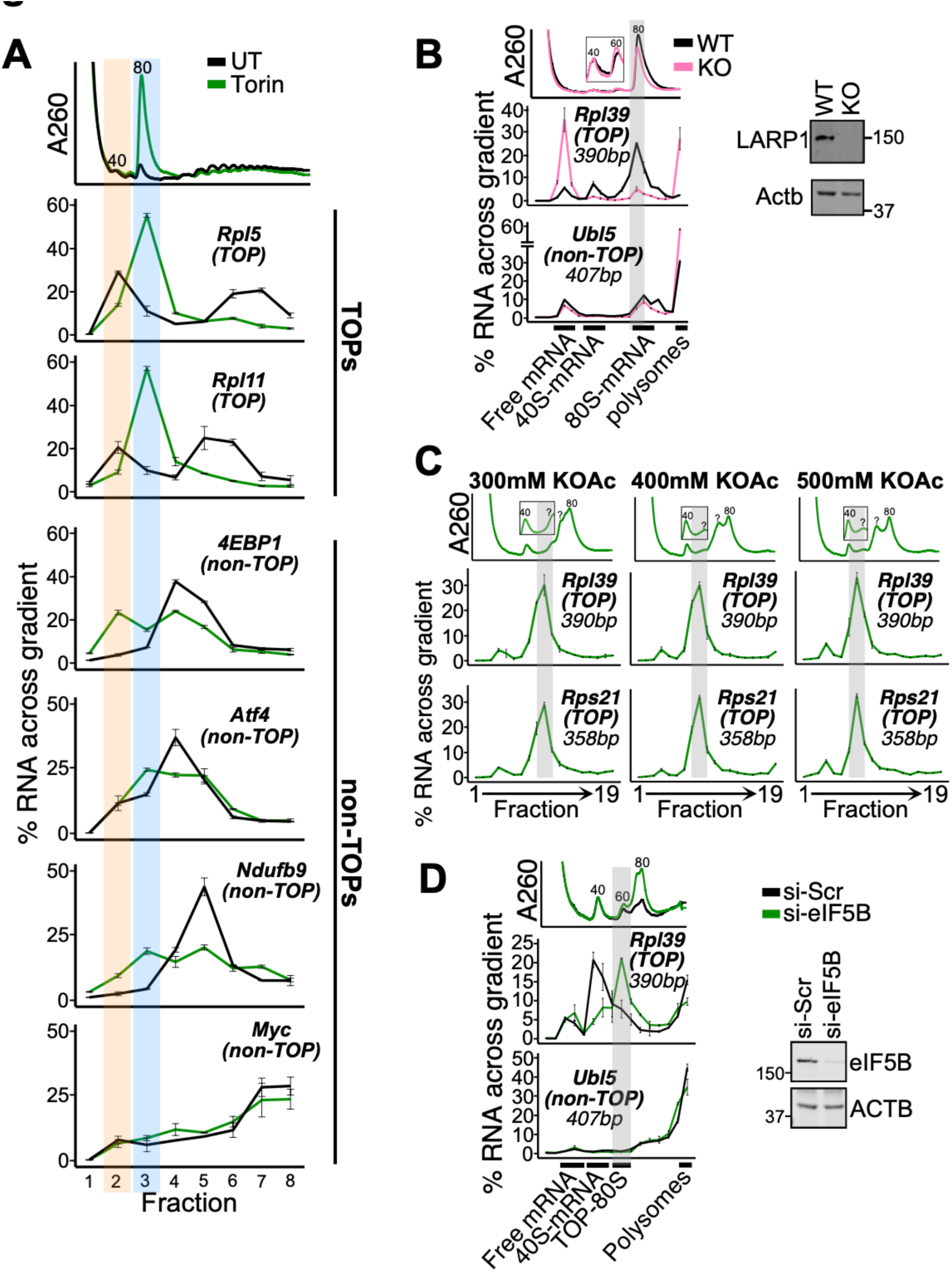
Characterization of the TOP-80S complex (related to **Fig. 1**). (**A**) qPCR for additional TOPs and non-TOPs from Fig. 1C. For clarity, the A260 trace is reproduced from Fig. 1C. (**B**) WT and LARP1-KO cells (HEK293T) were treated with 300 nM Torin1 for 1 h and lysates spread along 15-35% sucrose gradients containing 100 mM KOAc followed by qPCR against genes of interest. Western blot validating the LARP1-KO is shown. (**C**) HEK293T cells were treated with 300 nM Torin1 for 1 h and lysates fractionated along 15-35% sucrose gradients containing 300 mM, 400 mM, or 500 mM KOAc followed by qPCR against genes of interest. (**D**) HEK293T cells were treated with siRNAs targeting scrambled (Si-Scr) or eIF5B for 48 hours and lysates fractionated along 15-35% sucrose gradients containing 200 mM KOAc followed by qPCR against genes of interest. Western blot demonstrating knockdown efficiency is shown. For (A-D), data are shown from one experiment representative of two biological replicates (A) or one biological replicate (B-D). For qPCR data, error bars reflect the SD of 2-4 technical replicates from one experiment. Orange and blue highlights correspond to the TOP-40S and TOP-80S, respectively (A) and gray highlights correspond to the TOP-80S (B-D).

**Fig. S2.**
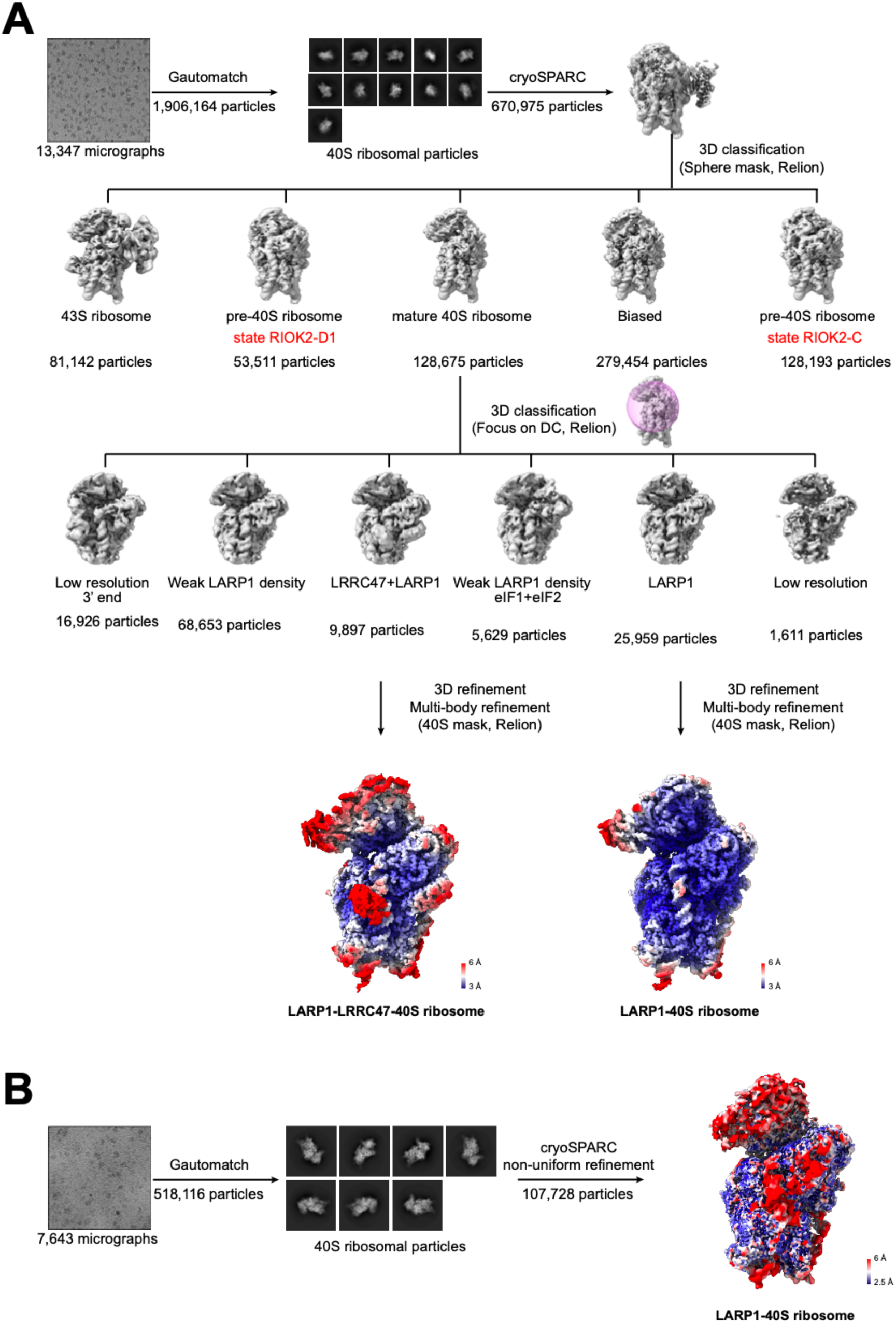
The sorting scheme of the cryo-EM dataset. (**A-B**) Sorting schemes of the cryo-EM datasets from the PYM1 (A) and LARP1 (B) immunoprecipitation samples. Masks and software used during data processing are shown. Maps are colored according to their local resolution distribution.

**Fig. S3.**
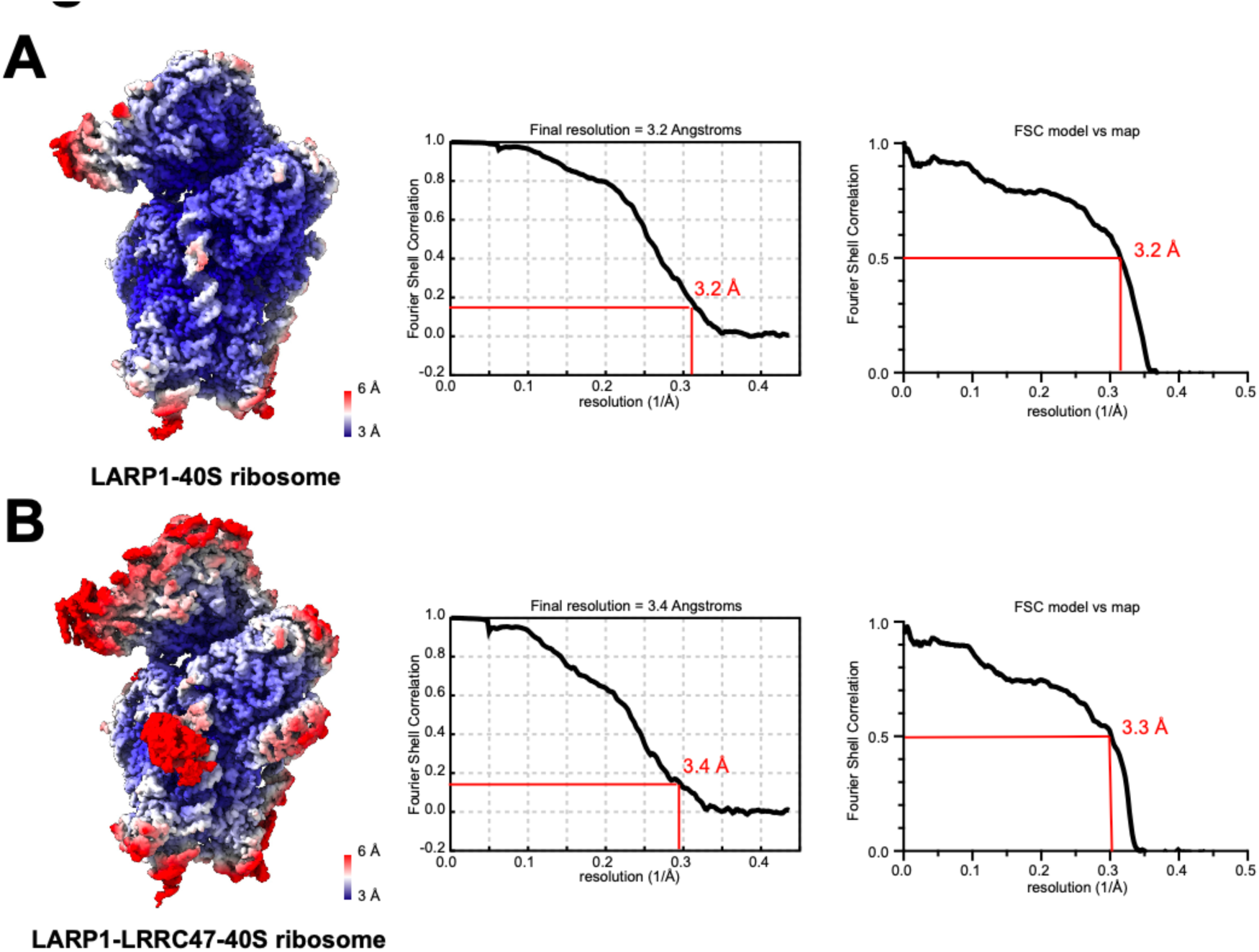
Local resolution and FSC curves of the LARP1-40S ribosome structures. (**A-B**) Cryo-EM maps of the LARP1-40S ribosome (A) and the LARP1-LRRC47-40S ribosome (B) are colored according to their local resolution estimation (left). The corresponding Fourier Shell Correlation (FSC) curves of the maps (middle) and model-to-map correlation curves (right) were calculated in Relion. The corresponding resolution was estimated using either the 0.143 or 0.5 cutoff criterion (red dotted line).

**Fig. S4.**
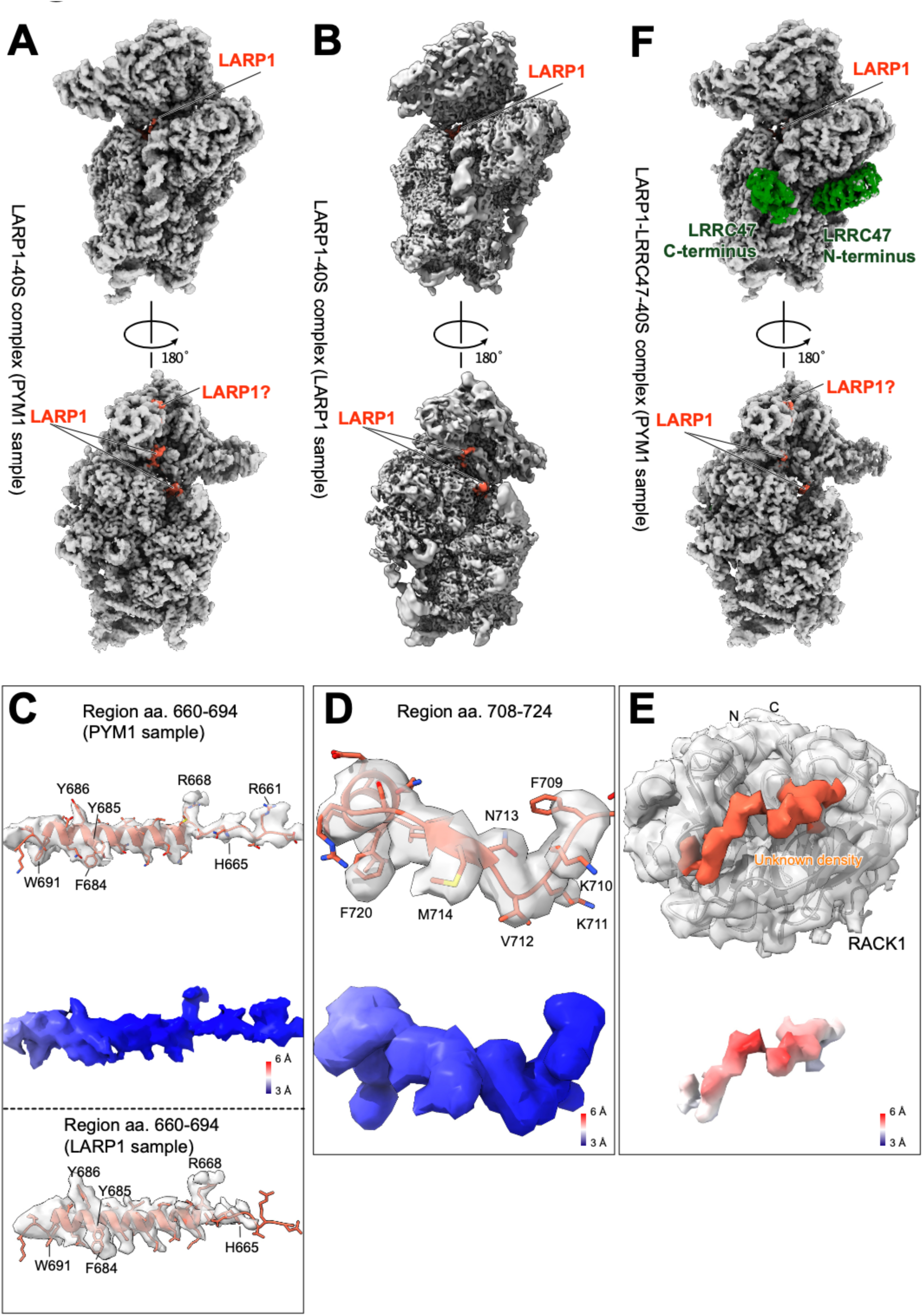
Detailed analysis of the interaction between LARP1 and the 40S ribosome. (**A-B**) Two different views of the density maps of the LARP1-40S ribosome structures obtained from the PYM1 sample (A) and the LARP1 sample (B). The maps are either filtered by DeepEMhancer (A) or cryoSPARC (B) according to their local resolution estimation. LARP1 (tomato red) is indicated. (**C-E**) Molecular models for LARP1 residues 660-694 (C), 708-724 (D), and the unknown region on RACK1 (E) are shown with density maps to support the assignment. The density map for region 660-694 is derived from the unsharpened map. Their relative maps are colored according to the local resolution estimated by Relion. Region 660-694 from the structure obtained from the LARP1 sample is also shown (C, bottom). (**F**) Two different views of the density map of the LARP1-LRRC47-40S ribosome structure obtained from the PYM1 sample. LARP1 (tomato red) and LRRC47 (green) are indicated.

**Fig. S5.**
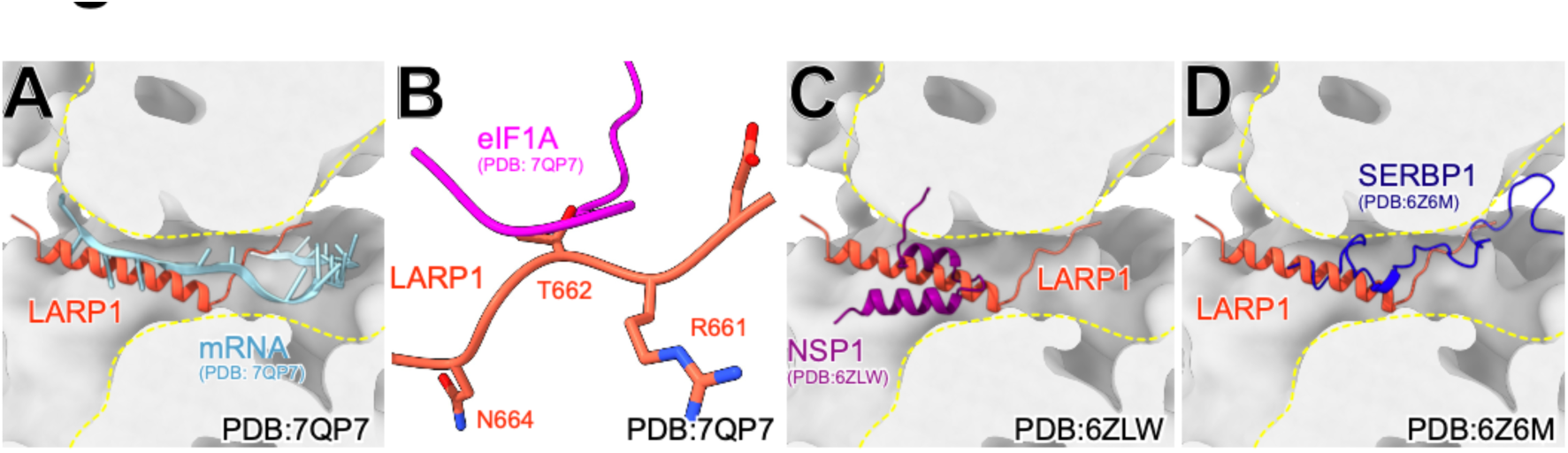
The LARP1 RBR inhibits mRNA binding. (**A-B**) Structural comparison of the LARP1-40S ribosome with the 48S initiation complex (PDB: 7QP7). The LARP1 RBR spatially clashes with mRNA in the mRNA channel (A) and with eIF1A near the DC (B). (**C-D**) Structural comparison of the LARP1 RBR with NSP1 (C) and SERBP1 (D) when bound in the mRNA channel.

**Fig. S6.**
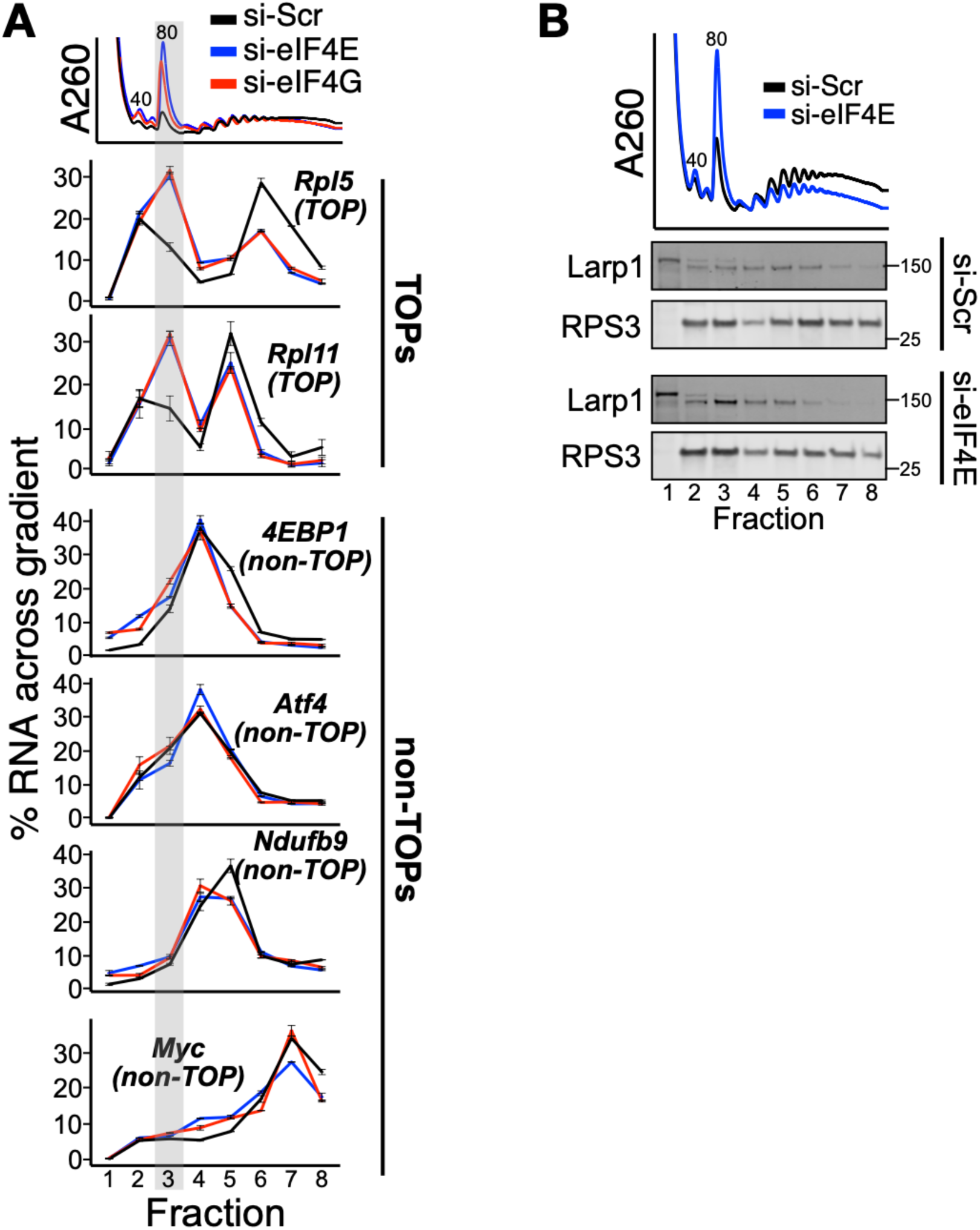
TOP mRNAs are repressed in 80S complexes by eIF4E or eIF4G knockdown. (**A**) qPCR for additional TOPs and non-TOPs from Fig. 4A. For clarity, the A260 trace is reproduced from Fig. 4A. Gray highlight corresponds to the TOP-80S. Data are shown from one experiment representative of two biological replicates. For qPCR data, error bars reflect the SD of 2-4 technical replicates from one experiment. (**B**) LARP1 sedimentation with the 80S ribosome is increased upon eIF4E knockdown. Protein was extracted from the gradient fractions and western blotted for LARP1 or RPS3.

**Fig. S7.**
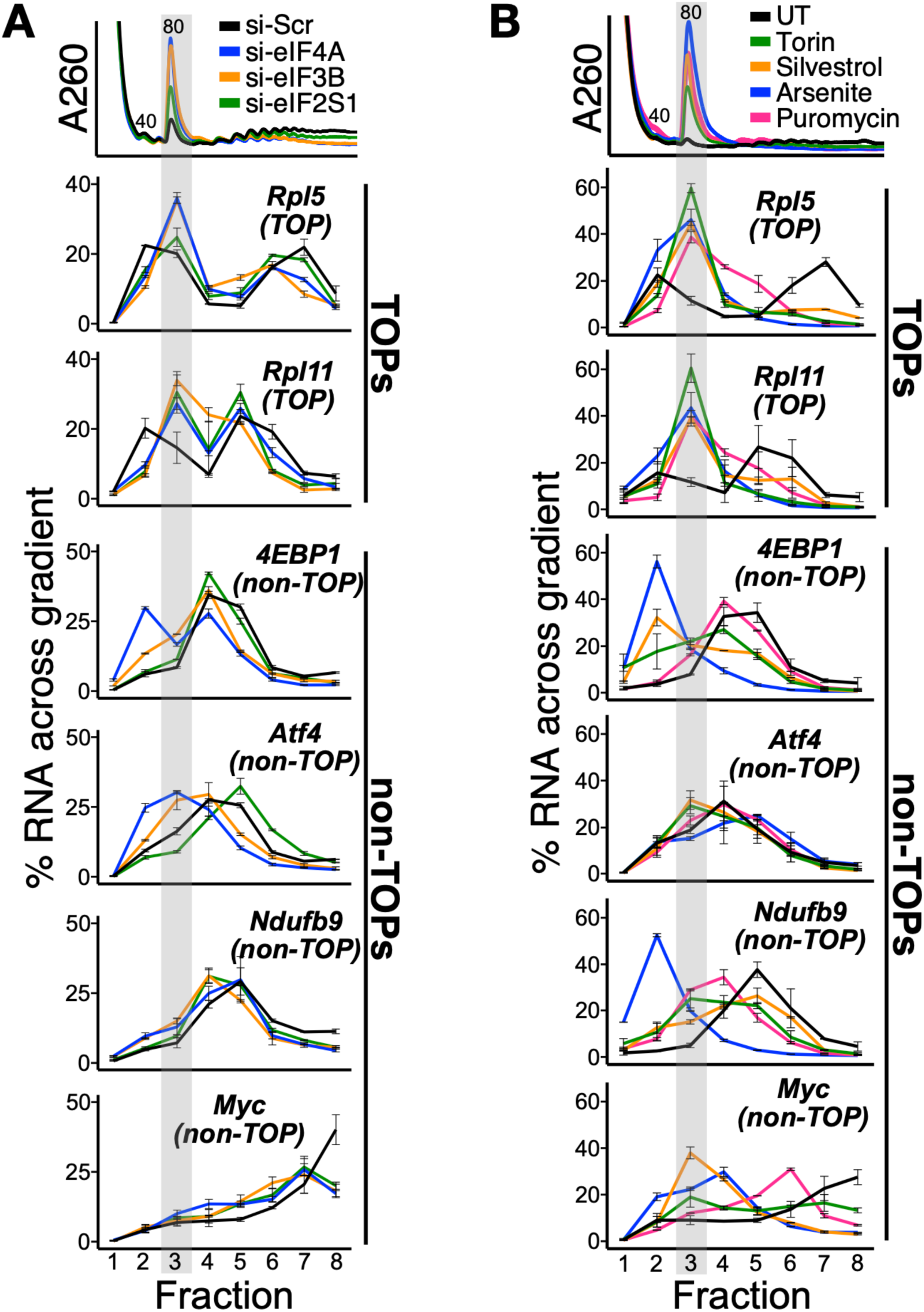
Free ribosomes induce TOP repression in 80S complexes. (**A-B**) qPCR for additional TOPs and non-TOPs from Fig. 4B (A) or Fig. 4D (B). For clarity, the A260 traces are reproduced from the respective main figures. Gray highlights correspond to the TOP-80S. Data are shown from one experiment representative of two biological replicates. For qPCR data, error bars reflect the SD of 2-4 technical replicates from one experiment.

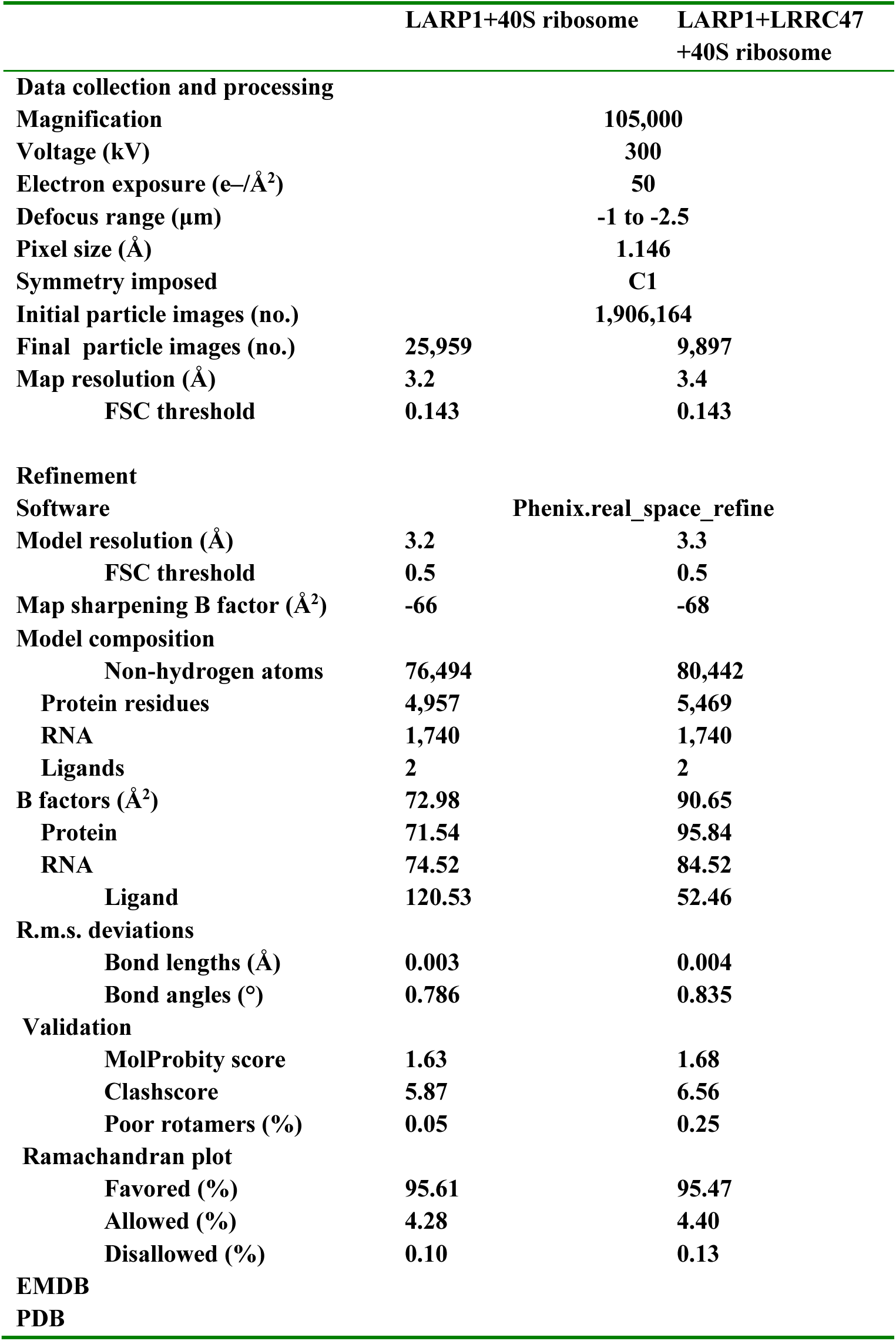
Table S1. Cryo-EM data statistics. Collection, refinement, and validation statistics are shown.

**Data S1. Mass spectrometry of the PYM1 sample.** LARP1 is a top hit among peptides identified from the PYM1 immunoprecipitation. Data table included as a supplemental file.

## References and Notes

1. O. Meyuhas, T. Kahan, The race to decipher the top secrets of TOP mRNAs. Biochim. Biophys. Acta. 1849, 801–811 (2015).

2. R. M. DePhilip, W. A. Rudert, I. Lieberman, Preferential stimulation of ribosomal protein synthesis by insulin and in the absence of ribosomal and messenger ribonucleic acid formation. Biochemistry. 19, 1662–1669 (1980).

3. P. K. Geyer, O. Meyuhas, R. P. Perry, L. F. Johnson, Regulation of ribosomal protein mRNA content and translation in growth-stimulated mouse fibroblasts. Mol. Cell. Biol. 2, 685–693 (1982).

4. O. Meyuhas, E. A. Thompson Jr, R. P. Perry, Glucocorticoids selectively inhibit translation of ribosomal protein mRNAs in P1798 lymphosarcoma cells. Mol. Cell. Biol. 7, 2691–2699 (1987).

5. R. Aloni, D. Peleg, O. Meyuhas, Selective translational control and nonspecific posttranscriptional regulation of ribosomal protein gene expression during development and regeneration of rat liver. Mol. Cell. Biol. 12, 2203–2212 (1992).

6. F. Loreni, G. Thomas, F. Amaldi, Transcription inhibitors stimulate translation of 5’ TOP mRNAs through activation of S6 kinase and the mTOR/FRAP signalling pathway. Eur. J. Biochem. 267, 6594–6601 (2000).

7. M. Stolovich, T. Lerer, Y. Bolkier, H. Cohen, O. Meyuhas, Lithium can relieve translational repression of TOP mRNAs elicited by various blocks along the cell cycle in a glycogen synthase kinase-3- and S6-kinase-independent manner. J. Biol. Chem. 280, 5336–5342 (2005).

8. W. J. Tuxworth Jr, H. Shiraishi, P. C. Moschella, K. Yamane, P. J. McDermott, D. Kuppuswamy, Translational activation of 5’-TOP mRNA in pressure overload myocardium. Basic Res. Cardiol. 103, 41–53 (2008).

9. C. Jouffe, G. Cretenet, L. Symul, E. Martin, F. Atger, F. Naef, F. Gachon, The circadian clock coordinates ribosome biogenesis. PLoS Biol. 11, e1001455 (2013).

10. R. Miloslavski, E. Cohen, A. Avraham, Y. Iluz, Z. Hayouka, J. Kasir, R. Mudhasani, S. N. Jones, N. Cybulski, M. A. Rüegg, O. Larsson, V. Gandin, A. Rajakumar, I. Topisirovic, O. Meyuhas, Oxygen sufficiency controls TOP mRNA translation via the TSC-Rheb-mTOR pathway in a 4E-BP-independent manner. J. Mol. Cell Biol. 6, 255–266 (2014).

11. K. A. Cottrell, R. C. Chiou, J. D. Weber, Upregulation of 5’-terminal oligopyrimidine mRNA translation upon loss of the ARF tumor suppressor. Sci. Rep. 10, 22276 (2020).

12. S. Levy, D. Avni, N. Hariharan, R. P. Perry, O. Meyuhas, Oligopyrimidine tract at the 5’ end of mammalian ribosomal protein mRNAs is required for their translational control. Proceedings of the National Academy of Sciences. 88 (1991), pp. 3319–3323.

13. M. L. Hammond, W. Merrick, L. H. Bowman, Sequences mediating the translation of mouse S16 ribosomal protein mRNA during myoblast differentiation and in vitro and possible control points for the in vitro translation. Genes Dev. 5, 1723–1736 (1991).

14. Y. Biberman, O. Meyuhas, Substitution of just five nucleotides at and around the transcription start site of rat beta-actin promoter is sufficient to render the resulting transcript a subject for translational control. FEBS Lett. 405, 333–336 (1997).

15. C. Bousquet-Antonelli, J.-M. Deragon, A comprehensive analysis of the La-motif protein superfamily. RNA. 15, 750–764 (2009).

16. B. D. Fonseca, C. Zakaria, J.-J. Jia, T. E. Graber, Y. Svitkin, S. Tahmasebi, D. Healy, H.-D. Hoang, J. M. Jensen, I. T. Diao, A. Lussier, C. Dajadian, N. Padmanabhan, W. Wang, E. Matta-Camacho, J. Hearnden, E. M. Smith, Y. Tsukumo, A. Yanagiya, M. Morita, E. Petroulakis, J. L. González, G. Hernández, T. Alain, C. K. Damgaard, La-related Protein 1 (LARP1) Represses Terminal Oligopyrimidine (TOP) mRNA Translation Downstream of mTOR Complex 1 (mTORC1). J. Biol. Chem. 290, 15996–16020 (2015).

17. R. M. Lahr, S. M. Mack, A. Héroux, S. P. Blagden, C. Bousquet-Antonelli, J.-M. Deragon, A. J. Berman, The La-related protein 1-specific domain repurposes HEAT-like repeats to directly bind a 5′TOP sequence. Nucleic Acids Res. 43, 8077–8088 (2015).

18. R. M. Lahr, B. D. Fonseca, G. E. Ciotti, H. A. Al-Ashtal, J.-J. Jia, M. R. Niklaus, S. P. Blagden, T. Alain, A. J. Berman, La-related protein 1 (LARP1) binds the mRNA cap, blocking eIF4F assembly on TOP mRNAs. Elife. 6, e24146 (2017).

19. L. Philippe, J.-J. Vasseur, F. Debart, C. C. Thoreen, La-related protein 1 (LARP1) repression of TOP mRNA translation is mediated through its cap-binding domain and controlled by an adjacent regulatory region. Nucleic Acids Res. 46, 1457–1469 (2018).

20. L. Philippe, A. M. G. van den Elzen, M. J. Watson, C. C. Thoreen, Global analysis of LARP1 translation targets reveals tunable and dynamic features of 5’ TOP motifs. Proc. Natl. Acad. Sci. U. S. A. 117, 5319–5328 (2020).

21. P. Fuentes, J. Pelletier, C. Martinez-Herráez, V. Diez-Obrero, F. Iannizzotto, T. Rubio, M. Garcia-Cajide, S. Menoyo, V. Moreno, R. Salazar, A. Tauler, A. Gentilella, The 40S-LARP1 complex reprograms the cellular translatome upon mTOR inhibition to preserve the protein synthetic capacity. Sci Adv. 7, eabg9275 (2021).

22. K. Ogami, Y. Oishi, K. Sakamoto, M. Okumura, R. Yamagishi, T. Inoue, M. Hibino, T. Nogimori, N. Yamaguchi, K. Furutachi, N. Hosoda, H. Inagaki, S.-I. Hoshino, mTOR- and LARP1-dependent regulation of TOP mRNA poly(A) tail and ribosome loading. Cell Rep. 41, 111548 (2022).

23. I. Patursky-Polischuk, M. Stolovich-Rain, M. Hausner-Hanochi, J. Kasir, N. Cybulski, J. Avruch, M. A. Rüegg, M. N. Hall, O. Meyuhas, The TSC-mTOR Pathway Mediates Translational Activation of TOP mRNAs by Insulin Largely in a Raptor- or Rictor-Independent Manner. Mol. Cell. Biol. 29, 1670 (2009).

24. B. B. Li, C. Qian, P. A. Gameiro, C.-C. Liu, T. Jiang, T. M. Roberts, K. Struhl, J. J. Zhao, Targeted profiling of RNA translation reveals mTOR-4EBP1/2-independent translation regulation of mRNAs encoding ribosomal proteins. Proceedings of the National Academy of Sciences. 115, E9325–E9332 (2018).

25. K. Haneke, J. Schott, D. Lindner, A. K. Hollensen, C. K. Damgaard, C. Mongis, M. Knop, W. Palm, A. Ruggieri, G. Stoecklin, CDK1 couples proliferation with protein synthesis. J. Cell Biol. 219 (2020), doi:10.1083/jcb.201906147.

26. H. B. Jefferies, C. Reinhard, S. C. Kozma, G. Thomas, Rapamycin selectively represses translation of the “polypyrimidine tract” mRNA family. Proc. Natl. Acad. Sci. U. S. A. 91, 4441–4445 (1994).

27. N. Terada, H. R. Patel, K. Takase, K. Kohno, A. C. Nairn, E. W. Gelfand, Rapamycin selectively inhibits translation of mRNAs encoding elongation factors and ribosomal proteins. Proc. Natl. Acad. Sci. U. S. A. 91, 11477–11481 (1994).

28. C. C. Thoreen, L. Chantranupong, H. R. Keys, T. Wang, N. S. Gray, D. M. Sabatini, A unifying model for mTORC1-mediated regulation of mRNA translation. Nature. 485 (2012), pp. 109–113.

29. Y. Yu, S.-O. Yoon, G. Poulogiannis, Q. Yang, X. M. Ma, J. Villén, N. Kubica, G. R. Hoffman, L. C. Cantley, S. P. Gygi, J. Blenis, Phosphoproteomic Analysis Identifies Grb10 as an mTORC1 Substrate That Negatively Regulates Insulin Signaling. Science. 332, 1322–1326 (2011).

30. S. A. Kang, M. E. Pacold, C. L. Cervantes, D. Lim, H. J. Lou, K. Ottina, N. S. Gray, B. E. Turk, M. B. Yaffe, D. M. Sabatini, mTORC1 phosphorylation sites encode their sensitivity to starvation and rapamycin. Science. 341, 1236566 (2013).

31. S. Hong, M. A. Freeberg, T. Han, A. Kamath, Y. Yao, T. Fukuda, T. Suzuki, J. K. Kim, K. Inoki, LARP1 functions as a molecular switch for mTORC1-mediated translation of an essential class of mRNAs. Elife. 6 (2017), doi:10.7554/eLife.25237.

32. B. D. Fonseca, R. M. Lahr, C. K. Damgaard, T. Alain, A. J. Berman, LARP1 on TOP of ribosome production. Wiley Interdiscip. Rev. RNA. 9, e1480 (2018).

33. J.-J. Jia, R. M. Lahr, M. T. Solgaard, B. J. Moraes, R. Pointet, A.-D. Yang, G. Celucci, T. E. Graber, H.-D. Hoang, M. R. Niklaus, I. A. Pena, A. K. Hollensen, E. M. Smith, M. Chaker-Margot, L. Anton, C. Dajadian, M. Livingstone, J. Hearnden, X.-D. Wang, Y. Yu, T. Maier, C. K. Damgaard, A. J. Berman, T. Alain, B. D. Fonseca, mTORC1 promotes TOP mRNA translation through site-specific phosphorylation of LARP1. Nucleic Acids Res. 49, 3461– 3489 (2021).

34. A. Gentilella, F. D. Morón-Duran, P. Fuentes, G. Zweig-Rocha, F. Riaño-Canalias, J. Pelletier, M. Ruiz, G. Turón, J. Castaño, A. Tauler, C. Bueno, P. Menéndez, S. C. Kozma, G. Thomas, Autogenous Control of 5′TOP mRNA Stability by 40S Ribosomes. Mol. Cell. 67, 55–70.e4 (2017).

35. C. Schneider, F. Erhard, B. Binotti, A. Buchberger, J. Vogel, U. Fischer, An unusual mode of baseline translation adjusts cellular protein synthesis capacity to metabolic needs. Cell Rep. 41, 111467 (2022).

36. E. Wolin, J. K. Guo, M. R. Blanco, A. A. Perez, I. N. Goronzy, A. A. Abdou, D. Gorhe, M. Guttman, M. Jovanovic, SPIDR: a highly multiplexed method for mapping RNA-protein interactions uncovers a potential mechanism for selective translational suppression upon cellular stress. bioRxiv (2023), doi:10.1101/2023.06.05.543769.

37. C. C. Thoreen, S. A. Kang, J. W. Chang, Q. Liu, J. Zhang, Y. Gao, L. J. Reichling, T. Sim, D. M. Sabatini, N. S. Gray, An ATP-competitive mammalian target of rapamycin inhibitor reveals rapamycin-resistant functions of mTORC1. J. Biol. Chem. 284, 8023–8032 (2009).

38. A. A. Infante, R. Baierlein, Pressure-induced dissociation of sedimenting ribosomes: effect on sedimentation patterns. Proc. Natl. Acad. Sci. U. S. A. 68, 1780–1785 (1971).

39. R. J. Beller, N. H. Lubsen, Effect of polypeptide chain length on dissociation of ribosomal complexes. Biochemistry. 11, 3271–3276 (1972).

40. M. Noll, B. Hapke, M. H. Schreier, H. Noll, Structural dynamics of bacterial ribosomes. I. Characterization of vacant couples and their relation to complexed ribosomes. J. Mol. Biol. 75, 281–294 (1973).

41. J. H. Lee, T. V. Pestova, B.-S. Shin, C. Cao, S. K. Choi, T. E. Dever, Initiation factor eIF5B catalyzes second GTP-dependent step in eukaryotic translation initiation. Proceedings of the National Academy of Sciences. 99, 16689–16694 (2002).

42. M. D. Diem, C. C. Chan, I. Younis, G. Dreyfuss, PYM binds the cytoplasmic exon-junction complex and ribosomes to enhance translation of spliced mRNAs. Nat. Struct. Mol. Biol. 14, 1173–1179 (2007).

43. J. Jumper, R. Evans, A. Pritzel, T. Green, M. Figurnov, O. Ronneberger, K. Tunyasuvunakool, R. Bates, A. Žídek, A. Potapenko, A. Bridgland, C. Meyer, S. A. A. Kohl, A. J. Ballard, A. Cowie, B. Romera-Paredes, S. Nikolov, R. Jain, J. Adler, T. Back, S. Petersen, D. Reiman, E. Clancy, M. Zielinski, M. Steinegger, M. Pacholska, T. Berghammer, S. Bodenstein, D. Silver, O. Vinyals, A. W. Senior, K. Kavukcuoglu, P. Kohli, D. Hassabis, Highly accurate protein structure prediction with AlphaFold. Nature. 596, 583– 589 (2021).

44. H. Schwenzer, M. Abdel Mouti, P. Neubert, J. Morris, J. Stockton, S. Bonham, M. Fellermeyer, J. Chettle, R. Fischer, A. D. Beggs, S. P. Blagden, LARP1 isoform expression in human cancer cell lines. RNA Biol. 18, 237–247 (2021).

45. S. Mattijssen, G. Kozlov, S. Gaidamakov, A. Ranjan, B. D. Fonseca, K. Gehring, R. J. Maraia, The isolated La-module of LARP1 mediates 3’ poly(A) protection and mRNA stabilization, dependent on its intrinsic PAM2 binding to PABPC1. RNA Biol. 18, 275–289 (2021).

46. G. Kozlov, S. Mattijssen, J. Jiang, S. Nyandwi, T. Sprules, J. R. Iben, S. L. Coon, S. Gaidamakov, A. M. Noronha, C. J. Wilds, R. J. Maraia, K. Gehring, Structural basis of 3’- end poly(A) RNA recognition by LARP1. Nucleic Acids Res. 50, 9534–9547 (2022).

47. H. A. Al-Ashtal, C. M. Rubottom, T. C. Leeper, A. J. Berman, The LARP1 La-Module recognizes both ends of TOP mRNAs. RNA Biol. 18, 248–258 (2021).

48. L. B. Jenner, N. Demeshkina, G. Yusupova, M. Yusupov, Structural aspects of messenger RNA reading frame maintenance by the ribosome. Nat. Struct. Mol. Biol. 17, 555–560 (2010).

49. J. Brito Querido, M. Sokabe, S. Kraatz, Y. Gordiyenko, J. M. Skehel, C. S. Fraser, V. Ramakrishnan, Structure of a human 48S translational initiation complex. Science. 369, 1220–1227 (2020).

50. S.-H. Yi, V. Petrychenko, J. E. Schliep, A. Goyal, A. Linden, A. Chari, H. Urlaub, H. Stark, M. V. Rodnina, S. Adio, N. Fischer, Conformational rearrangements upon start codon recognition in human 48S translation initiation complex. Nucleic Acids Res. 50, 5282–5298 (2022).

51. M. Thoms, R. Buschauer, M. Ameismeier, L. Koepke, T. Denk, M. Hirschenberger, H. Kratzat, M. Hayn, T. Mackens-Kiani, J. Cheng, J. H. Straub, C. M. Stürzel, T. Fröhlich, O. Berninghausen, T. Becker, F. Kirchhoff, K. M. J. Sparrer, R. Beckmann, Structural basis for translational shutdown and immune evasion by the Nsp1 protein of SARS-CoV-2. Science. 369, 1249–1255 (2020).

52. J. N. Wells, R. Buschauer, T. Mackens-Kiani, K. Best, H. Kratzat, O. Berninghausen, T. Becker, W. Gilbert, J. Cheng, R. Beckmann, Structure and function of yeast Lso2 and human CCDC124 bound to hibernating ribosomes. PLoS Biol. 18, e3000780 (2020).

53. M. Ameismeier, I. Zemp, J. van den Heuvel, M. Thoms, O. Berninghausen, U. Kutay, R. Beckmann, Structural basis for the final steps of human 40S ribosome maturation. Nature. 587, 683–687 (2020).

54. H. Kratzat, T. Mackens-Kiani, M. Ameismeier, M. Potocnjak, J. Cheng, E. Dacheux, A. Namane, O. Berninghausen, F. Herzog, M. Fromont-Racine, T. Becker, R. Beckmann, A structural inventory of native ribosomal ABCE1-43S pre-initiation complexes. EMBO J. 40, e105179 (2021).

55. Y. Huo, V. Iadevaia, Z. Yao, I. Kelly, S. Cosulich, S. Guichard, L. J. Foster, C. G. Proud, Stable isotope-labelling analysis of the impact of inhibition of the mammalian target of rapamycin on protein synthesis. Biochem. J. 444, 141–151 (2012).

56. E. M. Smith, N. E. H. Benbahouche, K. Morris, A. Wilczynska, S. Gillen, T. Schmidt, H. A. Meijer, R. Jukes-Jones, K. Cain, C. Jones, M. Stoneley, J. A. Waldron, C. Bell, B. D. Fonseca, S. Blagden, A. E. Willis, M. Bushell, The mTOR regulated RNA-binding protein LARP1 requires PABPC1 for guided mRNA interaction. Nucleic Acids Res. 49, 458–478 (2020).

57. T. Hochstoeger, P. Papasaikas, E. Piskadlo, J. A. Chao, Distinct roles of LARP1 and 4EBP1/2 in regulating translation and stability of 5′TOP mRNAs. bioRxiv (2023), p. 2023.05.22.541712.

58. A. Haghighat, S. Mader, A. Pause, N. Sonenberg, Repression of cap-dependent translation by 4E-binding protein 1: competition with p220 for binding to eukaryotic initiation factor-4E. EMBO J. 14, 5701–5709 (1995).

59. K. Pakos-Zebrucka, I. Koryga, K. Mnich, M. Ljujic, A. Samali, A. M. Gorman, The integrated stress response. EMBO Rep. 17, 1374–1395 (2016).

60. A. Haghighat, N. Sonenberg, eIF4G dramatically enhances the binding of eIF4E to the mRNA 5’-cap structure. J. Biol. Chem. 272, 21677–21680 (1997).

61. A. Yanagiya, Y. V. Svitkin, S. Shibata, S. Mikami, H. Imataka, N. Sonenberg, Requirement of RNA binding of mammalian eukaryotic translation initiation factor 4GI (eIF4GI) for efficient interaction of eIF4E with the mRNA cap. Mol. Cell. Biol. 29, 1661–1669 (2009).

62. I. A. Nikonorova, E. T. Mirek, C. C. Signore, M. P. Goudie, R. C. Wek, T. G. Anthony, Time-resolved analysis of amino acid stress identifies eIF2 phosphorylation as necessary to inhibit mTORC1 activity in liver. J. Biol. Chem. 293, 5005–5015 (2018).

63. J. Averous, S. Lambert-Langlais, F. Mesclon, V. Carraro, L. Parry, C. Jousse, A. Bruhat, A.-C. Maurin, P. Pierre, C. G. Proud, P. Fafournoux, GCN2 contributes to mTORC1 inhibition by leucine deprivation through an ATF4 independent mechanism. Sci. Rep. 6, 27698 (2016).

64. A. L. Wolfe, K. Singh, Y. Zhong, P. Drewe, V. K. Rajasekhar, V. R. Sanghvi, K. J. Mavrakis, M. Jiang, J. E. Roderick, J. Van der Meulen, J. H. Schatz, C. M. Rodrigo, C. Zhao, P. Rondou, E. de Stanchina, J. Teruya-Feldstein, M. A. Kelliher, F. Speleman, J. A. Porco Jr, J. Pelletier, G. Rätsch, H.-G. Wendel, RNA G-quadruplexes cause eIF4A-dependent oncogene translation in cancer. Nature. 513, 65–70 (2014).

65. R. F. Duncan, J. W. Hershey, Translational repression by chemical inducers of the stress response occurs by different pathways. Arch. Biochem. Biophys. 256, 651–661 (1987).

66. M. E. Azzam, I. D. Algranati, Mechanism of Puromycin Action: Fate of Ribosomes after Release of Nascent Protein Chains from Polysomes. Proceedings of the National Academy of Sciences. 70, 3866–3869 (1973).

67. J. R. Wiśniewski, M. Y. Hein, J. Cox, M. Mann, A “proteomic ruler” for protein copy number and concentration estimation without spike-in standards. Mol. Cell. Proteomics. 13, 3497– 3506 (2014).

68. V. E. Velculescu, S. L. Madden, L. Zhang, A. E. Lash, J. Yu, C. Rago, A. Lal, C. J. Wang, G. A. Beaudry, K. M. Ciriello, B. P. Cook, M. R. Dufault, A. T. Ferguson, Y. Gao, T. C. He, H. Hermeking, S. K. Hiraldo, P. M. Hwang, M. A. Lopez, H. F. Luderer, B. Mathews, J. M. Petroziello, K. Polyak, L. Zawel, K. W. Kinzler, Analysis of human transcriptomes. Nat. Genet. 23, 387–388 (1999).

69. R. Duncan, J. W. Hershey, Identification and quantitation of levels of protein synthesis initiation factors in crude HeLa cell lysates by two-dimensional polyacrylamide gel electrophoresis. J. Biol. Chem. 258, 7228–7235 (1983).

70. K. Aoki, S. Adachi, M. Homoto, H. Kusano, K. Koike, T. Natsume, LARP1 specifically recognizes the 3’ terminus of poly(A) mRNA. FEBS Lett. 587, 2173–2178 (2013).

71. J. A. Saba, K. Liakath-Ali, R. Green, F. M. Watt, Translational control of stem cell function. Nat. Rev. Mol. Cell Biol. 22, 671–690 (2021).

72. M. Delarue, G. P. Brittingham, S. Pfeffer, I. V. Surovtsev, S. Pinglay, K. J. Kennedy, M. Schaffer, J. I. Gutierrez, D. Sang, G. Poterewicz, J. K. Chung, J. M. Plitzko, J. T. Groves, C. Jacobs-Wagner, B. D. Engel, L. J. Holt, mTORC1 Controls Phase Separation and the Biophysical Properties of the Cytoplasm by Tuning Crowding. Cell. 174, 338–349.e20 (2018).

73. D. C. Fingar, S. Salama, C. Tsou, E. Harlow, J. Blenis, Mammalian cell size is controlled by mTOR and its downstream targets S6K1 and 4EBP1/eIF4E. Genes Dev. 16, 1472–1487 (2002).

74. R. K. Khajuria, M. Munschauer, J. C. Ulirsch, C. Fiorini, L. S. Ludwig, S. K. McFarland, N. J. Abdulhay, H. Specht, H. Keshishian, D. R. Mani, M. Jovanovic, S. R. Ellis, C. P. Fulco, J. M. Engreitz, S. Schütz, J. Lian, K. W. Gripp, O. K. Weinberg, G. S. Pinkus, L. Gehrke, A. Regev, E. S. Lander, H. T. Gazda, W. Y. Lee, V. G. Panse, S. A. Carr, V. G. Sankaran, Ribosome Levels Selectively Regulate Translation and Lineage Commitment in Human Hematopoiesis. Cell. 173, 90–103.e19 (2018).

75. E. W. Mills, R. Green, N. T. Ingolia, Slowed decay of mRNAs enhances platelet specific translation. Blood. 129 (2017), pp. e38–e48.

76. M. Mura, T. G. Hopkins, T. Michael, N. Abd-Latip, J. Weir, E. Aboagye, F. Mauri, C. Jameson, J. Sturge, H. Gabra, M. Bushell, A. E. Willis, E. Curry, S. P. Blagden, LARP1 post-transcriptionally regulates mTOR and contributes to cancer progression. Oncogene. 34, 5025–5036 (2015).

77. Z. Xu, J. Xu, H. Lu, B. Lin, S. Cai, J. Guo, F. Zang, R. Chen, LARP1 is regulated by the XIST/miR-374a axis and functions as an oncogene in non-small cell lung carcinoma. Oncol. Rep. 38, 3659–3667 (2017).

78. C. Xie, L. Huang, S. Xie, D. Xie, G. Zhang, P. Wang, L. Peng, Z. Gao, LARP1 predict the prognosis for early-stage and AFP-normal hepatocellular carcinoma. J. Transl. Med. 11, 272 (2013).

79. L. Ye, S.-T. Lin, Y.-S. Mi, Y. Liu, Y. Ma, H.-M. Sun, Z.-H. Peng, J.-W. Fan, Overexpression of LARP1 predicts poor prognosis of colorectal cancer and is expected to be a potential therapeutic target. Tumour Biol. 37, 14585–14594 (2016).

80. T. G. Hopkins, M. Mura, H. A. Al-Ashtal, R. M. Lahr, N. Abd-Latip, K. Sweeney, H. Lu, J. Weir, M. El-Bahrawy, J. H. Steel, S. Ghaem-Maghami, E. O. Aboagye, A. J. Berman, S. P. Blagden, The RNA-binding protein LARP1 is a post-transcriptional regulator of survival and tumorigenesis in ovarian cancer. Nucleic Acids Res. 44, 1227–1246 (2016).

81. M. Ameismeier, J. Cheng, O. Berninghausen, R. Beckmann, Visualizing late states of human 40S ribosomal subunit maturation. Nature. 558, 249–253 (2018).

82. I. Zemp, T. Wild, M.-F. O’Donohue, F. Wandrey, B. Widmann, P.-E. Gleizes, U. Kutay, Distinct cytoplasmic maturation steps of 40S ribosomal subunit precursors require hRio2. J. Cell Biol. 185, 1167–1180 (2009).

83. S. Q. Zheng, E. Palovcak, J.-P. Armache, K. A. Verba, Y. Cheng, D. A. Agard, MotionCor2: anisotropic correction of beam-induced motion for improved cryo-electron microscopy. Nat. Methods. 14, 331–332 (2017).

84. K. Zhang, Gctf: Real-time CTF determination and correction. J. Struct. Biol. 193, 1–12 (2016).

85. J. Zivanov, T. Nakane, B. O. Forsberg, D. Kimanius, W. J. Hagen, E. Lindahl, S. H. Scheres, New tools for automated high-resolution cryo-EM structure determination in RELION-3. Elife. 7 (2018), doi:10.7554/eLife.42166.

86. A. Punjani, J. L. Rubinstein, D. J. Fleet, M. A. Brubaker, cryoSPARC: algorithms for rapid unsupervised cryo-EM structure determination. Nat. Methods. 14, 290–296 (2017).

87. R. Sanchez-Garcia, J. Gomez-Blanco, A. Cuervo, J. M. Carazo, C. O. S. Sorzano, J. Vargas, DeepEMhancer: a deep learning solution for cryo-EM volume post-processing. Commun Biol. 4, 874 (2021).

88. P. Emsley, K. Cowtan, Coot: model-building tools for molecular graphics. Acta Crystallogr. D Biol. Crystallogr. 60, 2126–2132 (2004).

89. P. D. Adams, P. V. Afonine, G. Bunkóczi, V. B. Chen, I. W. Davis, N. Echols, J. J. Headd, L.-W. Hung, G. J. Kapral, R. W. Grosse-Kunstleve, A. J. McCoy, N. W. Moriarty, R. Oeffner, R. J. Read, D. C. Richardson, J. S. Richardson, T. C. Terwilliger, P. H. Zwart, PHENIX: a comprehensive Python-based system for macromolecular structure solution. Acta Crystallogr. D Biol. Crystallogr. 66, 213–221 (2010).

90. V. B. Chen, W. B. Arendall 3rd, J. J. Headd, D. A. Keedy, R. M. Immormino, G. J. Kapral, L. W. Murray, J. S. Richardson, D. C. Richardson, MolProbity: all-atom structure validation for macromolecular crystallography. Acta Crystallogr. D Biol. Crystallogr. 66, 12–21 (2010).

91. T. D. Goddard, C. C. Huang, E. C. Meng, E. F. Pettersen, G. S. Couch, J. H. Morris, T. E. Ferrin, UCSF ChimeraX: Meeting modern challenges in visualization and analysis. Protein Sci. 27, 14–25 (2018).

